# Highly efficient and super-bright neurocircuit tracing using vector mixing-based virus cocktail

**DOI:** 10.1101/705772

**Authors:** Pei Sun, Sen Jin, Sijue Tao, Junjun Wang, Anan Li, Ning Li, Yang Wu, Jianxia Kuang, Yurong Liu, Lizhao Wang, Huimin Lin, Xiaohua Lv, Xiuli Liu, Jundan Peng, Siyu Zhang, Min Xu, Zhen Luo, Xiaobin He, Tonghui Xu, Xiangning Li, Shaoqun Zeng, Yu-Hui Zhang, Fuqiang Xu

**Affiliations:** Britton Chance Center for Biomedical Photonics, Wuhan National Laboratory for Optoelectronics-Huazhong University of Science and Technology, Wuhan, Hubei 430074, China; Centre for Brain Science, State Key Laboratory of Magnetic Resonance and Atomic Molecular Physics, Key Laboratory of Magnetic Resonance in Biological Systems, Wuhan Institute of Physics and Mathematics, CAS Centre for Excellence in Brain Science and Intelligence Technology, Chinese Academy of Sciences, Wuhan 430071, China; MoE Key Laboratory for Biomedical Photonics, School of Engineering Sciences, Huazhong University of Science and Technology, Wuhan, Hubei 430074, China; Shenzhen Institutes of Advanced Technology, Chinese Academy of Sciences, Shenzhen 518055, China; Brain Cognition and Brain Disease Institute (BCBDI), Shenzhen Institutes of Advanced Technology, Chinese Academy of Sciences, Shenzhen-Hong Kong Institute of Brain Science-Shenzhen Fundamental Research Institutions, Shenzhen 518055, China; Institute of Neuroscience, CAS Center for Excellence in Brain Science and Intelligence Technology, Chinese Academy of Sciences, 200031, Shanghai, China; Department of Anatomy and Physiology, Shanghai Jiao Tong University School of Medicine, 200025, Shanghai, China

## Abstract

Mapping the detailed cell-type-specific input networks and neuronal projectomes are essential to understand brain function in normal and pathological states. However, several properties of current tracing systems, including labeling sensitivity, trans-synaptic efficiencies, reproducibility among different individuals and different Cre-driver animals, still remained unsatisfactory. Here, we developed MAP-ENVIVIDERS, a recombinase system-dependent vector mixing-based strategy for highly efficient neurocircuit tracing. MAP-ENVIVIDERS enhanced tracing efficiency of input networks across the whole brain, with over 10-fold improvement in diverse previously poor-labeled input brain regions and particularly, up to 70-fold enhancement in brainstem compared with the current standard rabies-virus-mediated systems. MAP-ENVIVIDERS was over 10-fold more sensitive for cell-type-specific labeling than previous strategies, enabling us to capture individual cell-type-specific neurons with extremely complex axonal branches and presynaptic axonal boutons, both about one order of magnitude than previously reported and considered. MAP-ENVIVIDERS provides powerful tools for deconstructing novel input/output circuitry towards functional studies and disorders-related mechanisms.

## INTRODUCTION

Efficient mapping the input-output circuits of specific cell types is a central goal in neuroscience. Comprehensive tracing presynaptic partners to specific cell types is essential to understand how the input resources and strength received from other brain regions and other type of neurons guided the corresponding behaviors. For cell-type-specific input mapping, currently, rabies virus (RV, SAD B19 strain)-mediated monosynaptic tracing systems with the involvement of two or more recombinant adeno-associated viruses (rAAVs) helpers have been widely used for tracing the direct input network of a given type of neuron^1–6^. However, problems such as underestimated total input numbers, inefficient trans-synaptic abilities, particularly leading to rare labeling in brain regions which were far away from injection site or with weaker input strength, still hindered our further comprehensive characterization of neural circuits^7–9^. To solve this problem, extensive attention has been made via modifying the genome, including engineering of promoters for enhanced regulation strength or improvement of monosynaptic tracing efficiency with optimized glycoprotein (oG) to improve the efficiency of rabies glycopreotein (RG)-expressing helper rAAVs used in current RV-mediated monosynaptic tracing systems^5, 10^. Though great achievements have been made with these modifications, strategies to achieve comprehensive improvement of tracing efficiencies across the whole brain, particularly in previously poor-labeled brain regions were still in urgent need.

On the other hand, obtaining detailed single-neuron projectomes at the whole brain level enhances our understanding of input-output relationship of neural circuits and facilitates identification of novel cell types^11^. Though great achievements have been made in full morphology reconstruction of individual neurons belonging to different cell types^11–14^, fewer studies have focused on synaptic levels which still remained a great challenge due to the requirement of extremely higher labeling brightness (generally 3-fold higher) to distinguish the tiny-sized boutons (1-2 μm in diameter) from their resided thinner axons^15, 16^. Remarkably, understanding detailed axonal projection patterns on synaptic levels facilitate the elucidation of how information flows across diverse brain regions, to what extent individual neurons exert effects on their postsynaptic targets and how synaptic plasticity affects function of individual neurons under the normal and pathological conditions^17, 18^. To achieve neuronal sparse labeling with cell-type specificity, current strategy ultilized combinatorial genetic (e.g. Cre-driver animals) and viral method (e.g. rAAVs) based on intersectional gene expressions^19^. In these strategies, cell-type specificity was controlled by Cre-driver animals, labeling sparseness and brightness were mainly controlled by the mixtures of two or more rAAVs expressing essential elements for recombination (e.g. Cre/lox-Flp/FRT or Cre/lox-tTA/TRE combinations) along with the strength of Cre reombinase^13, 20, 21^. However, due to inhomogeneous strength of Cre recombinase, these labeling systems suffered from varied labeling brightness among different transgenic lines and rather low gene expression levels particularly in transgenic lines with weaker strength of Cre recombinase^21–23^. Thus, to generate detailed single-neuron projectomes of diverse cell types on synaptic levels, the optimal cell-type-specific labeling strategies should be with adjustable sparseness along with achievement of superb-bright labeling in diverse Cre-driver transgenic lines.

Overall, almost all current neural circuit labeling strategy involving multiple rAAV vectors focused on modifying viral genomes to improve vector properties of each separately packaged rAAV. However, when used together via co-injections or co-administrations, the labeling efficiencies or brightness of these multiple rAAV vectors still remained unsatisfactory. The probable reason was that such individual, separate improvements failed to consider interactions among different rAAVs. In fact, previous studies indicated that mutual suppression is prevalent among different types of viruses, e.g., AAV over herpes simplex virus^24^, AAV over adenovirus^25^, or even among the same types of viruses (e.g., herpes simplex virus and pseudorabies virus)^26–28^. Taken these facts, we reasoned that mutual suppression between different rAAVs, though have not been previously reported (to the best of our knowledge), will also significantly affect the gene expression levels. We further reasoned that altering viral packaging strategy, i.e., replacing the package of each viral vector independently by mixing multiple vectors in a single step on cellular level (i.e., copackaging strategy)^29, 30^ would enhance multi-gene expression efficiencies in neural circuit studies.

Based on this hypothesis, we developed MAP-ENVIVIDERS (**m**ultifaceted **a**melioration **p**rocess to **e**nhance **n**eurocircuit **vi**sualization by **vi**ral vectors **de**pending on **r**ecombinase **s**ystems), a recombinase system-dependent viral copackaging-based strategy for highly efficient neural circuit tracing. With MAP-ENVIVIDERS, we achieved: (i) 10- to 70-fold increased tracing efficiencies in 40% of all input brain regions for trans-monosynaptic retrograde tracing with RV-mediated systems; (ii) More than an order of magnitude of labeling brightness for cell-type-specific labeling than previous strategies and unbiased, super-bright labeling among diverse transgenic mice. Combined with whole-brain imaging systems, MAP-ENVIVIDERS enables identification of novel projection patterns and capture presynaptic axonal boutons of individual cell-type-specific neurons with extreme complexity. Finally, we demonstrated that viral copackaging strategy significantly ameliorated mutual suppression among different rAAVs and enhanced compatibility of multi-gene expressions in the same cells, the probable reasons leading to the high sensitivity and efficiency of MAP-ENVIVIDERS.

## RESULTS

### MAP-ENVIVIDERS substantially improves the efficiency of RV-mediated monosynaptic tracing systems

The current RV-mediated trans-monosynaptic tracing systems involve two different processes: the entry of EnvA-pseudotyped, RG-deleted RV (EnvA-RVΔG) into the cells through the interaction of EnvA with TVA receptor delivered by one rAAV, and the assembly of infectious RV by providing RVΔG with the RG delivered by another rAAV^1–6^. To investigate whether MAP-ENVIVIDERS, the recombinase system-dependent viral copackaging strategy would improve the coherence between the TVA-EnvA interaction and RVΔG-RG assembly, and thus leading to a significant increase in the efficiency of trans-monosynaptic labeling with the prevalent rAAV-RV system, we designed MAP-ENVIVIDERS with two Cre-dependent vectors, AAV-DIO-EGFP-TVA (GT) and AAV-DIO-RG (**Fig. 1a**). As an example, we copackaged AAV-DIO-GT and AAV-DIO-RG at a ratio of 1:2 to generate littermate virus, termed lAAV-DIO-GT/RG for convenience (where l refers to littermate hereafter). For comparisons, we packaged the two vectors independently and mixed them at a ratio of 1:2, the mixtures were abbreviated for sAAV-DIO-GT/RG (s refers to stranger hereafter; **Fig. 1b**). Since genetically modified, EnvA-pseudotyped RV (EnvA-SADΔG-DsRed) used in two systems were identical, comparisons of tracing efficiencies between lAAV-DIO-GT/RG and sAAV-DIO-GT/RG represented the difference between MAP-ENVIVIDERS and prevalent rAAV-RV system.

**Fig. 1.**
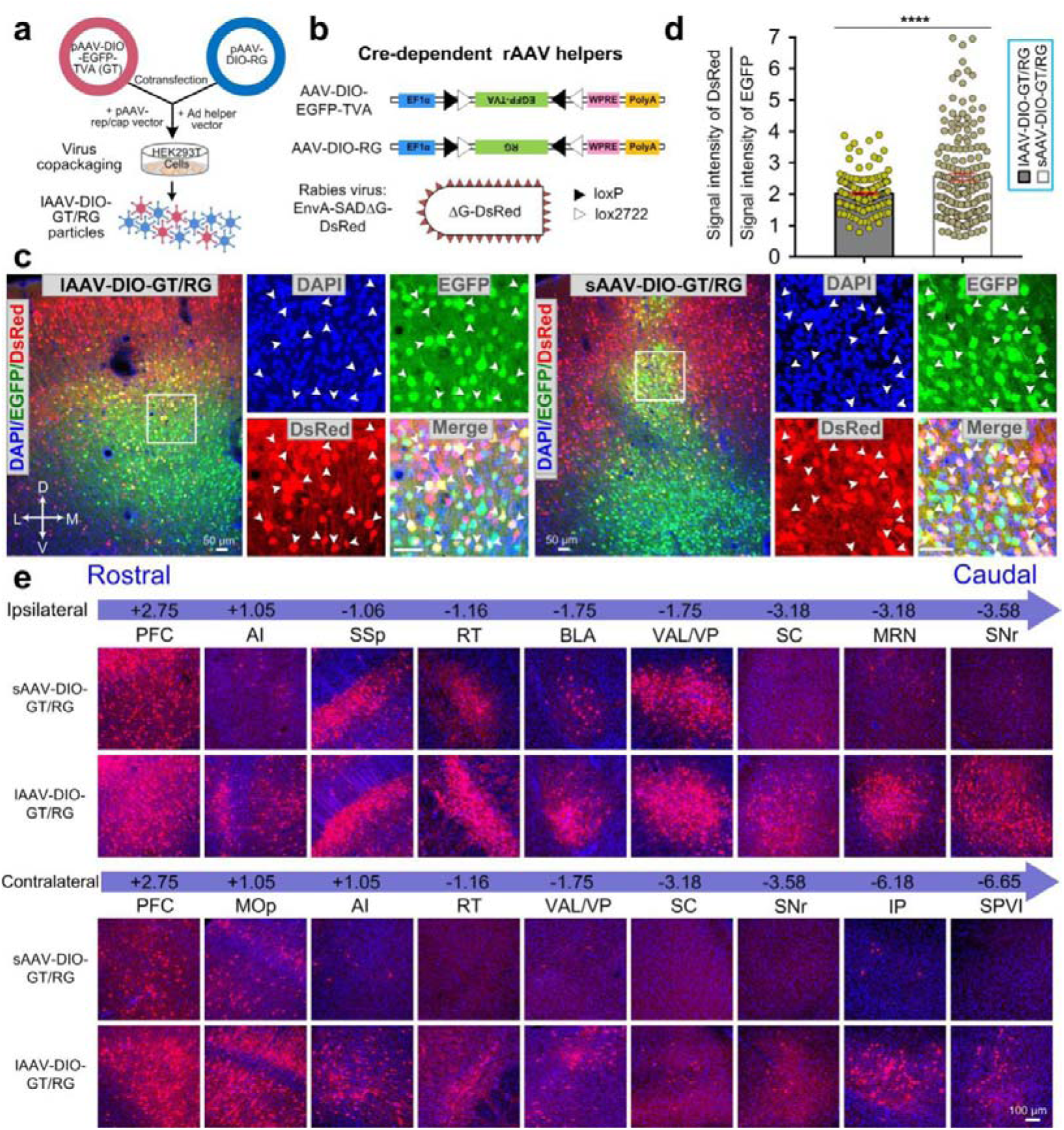
Demonstration of MAP-ENVIVDERS in tracing direct input networks in MOp. **a**, Schematic for generating copackaged rAAV helper by MAP-ENVIVDERS (lAAV-DIO-GT/RG). Briefly, two Cre-dependent AAV vectors for the respective expression of TVA and RG were premixed at a ratio of 1:2. **b**, Schematic of the two Cre-dependent rAAV helpers and the EnvA-pseudotyped rabies virus. **c**, Representative coronal sections showing the injection site in lAAV-DIO-GT/RG- and sAAV-DIO-GT/RG-labeled mice. Boxed regions showed colocalization of 4′,6-diamidino-2- phenylindole (DAPI, blue), EGFP (green), and DsRed (red). White arrowheads indicated triple-positive neurons. **d**, Comparison of the ratio between the signal intensity of DsRed and EGFP in starter cells labeled by lAAV-DIO-GT/RG and sAAV-DIO-GT/RG, n = 170 neurons from 3 mice for both groups. **e**, Representative images showing ipsi- and contralateral input neurons in different brain regions labeled by EnvA-pseudotyped rabies virus via lAAV-DIO-GT/RG or sAAV-DIO-GT/RG. Data are presented as mean ± s.e.m.; two-tailed t-test, ****P < 0.0001. Abbreviations: see **Supplementary Table 2**.

We injected equal volume of lAAV-DIO-GT/RG or sAAV-DIO-GT/RG into the primary motor cortex (MOp) of Thy1-Cre transgenic mice, followed by the injection of EnvA-pseudotyped RV 21 days later. Nine days after the second injection, the mice were sacrificed for analysis. We found that the numbers of EGFP^+^/DsRed^+^ starter cells in lAAV-DIO-GT/RG and sAAV-DIO-GT/RG groups were similar (2,386 ± 172 vs. 2,039 ± 127, n = 5 animals each; **Fig. 1c** and **Supplementary Table 1**) and the value of the DsRed/EGFP ratio^31^ closely matched the preset ratio of 2:1 for the lAAV-DIO-GT/RG group but significantly deviated from 2:1 for the sAAV-DIO-GT/RG group (**Fig. 1d** and **Supplementary Fig. 1a**, left panel; 2.01 ± 0.04 versus 2.67 ± 0.13; mean ± s.e.m., unpaired t-test, P < 0.0001, n = 3 mice). Furthermore, the distribution of ratios in individual neurons was significantly narrower for lAAV-DIO-GT/RG than sAAV-DIO-GT/RG (**Fig. 1d**). We next examined the inputs of two labeling groups qualitatively and fluorescent images showed that the improvement granted by the MAP-ENVIVIDERS system was most prominent for highly underrepresented regions far from the injection site, such as the reticular nucleus of the thalamus (RT), the superior colliculus (SC), the midbrain reticular nucleus (MRN) and the substantia nigra pars reticulata (SNr) in midbrain on the ipsilateral side and all regions on the contralateral side, especially the interposed nucleus (IP) in cerebellum and the spinal nucleus of the trigeminal nerve, subnucleus interpolaris (SPVI) in medulla. Furthermore, MAP-ENVIVIDERS also improved the labeling moderately in the brain regions that were well-labeled by the classical sAAV-DIO-GT/RG-RV combination, such as the prefrontal cortex (PFC), the primary somatosensory area (SSp), and the ventral anterior-lateral complex of the thalamus (VAL)/ventral posterior complex of the thalamus (VP) (**Fig. 1e** and **Supplementary Fig. 2,3**).

To compare the global tracing efficiency of the two systems quantitatively, we continued to divide the whole brain (+3.25 to −7.25 from the bregma) into 11 major regions containing 44 subregions according to the Allen Mouse Brain Atlas and registered the numbers of DsRed-labeled input cells in each subregion^32, 33^. We summarized the sparsely labeled input brain regions in sAAV-DIO-GT/RG labeling groups across the whole brain and found that number of input neurons in most of these areas were at least one order of magnitude enhanced by MAP-ENVIVIDERS (lAAV-DIO-GT/RG) (**Fig. 2a**). Such improvements covered diverse levels: (i) from hundreds to several thousands of neurons, such as visceral area (VISC) in cortical subplate (CTXsp), CP and ventral posteromedial nucleus of the thalamus (VPM); (ii) from dozens to several hundreds of neurons: such as central amygdalar nucleus (CEA) in striatum, ventral posterolateral nucleus of the thalamus (VPL), zona incerta (ZI) in hypothalamus, periaqueductal gray (PAG), pretectal region (PRT) and SNr in midbrain; (iii) from dozens to more than one thousands of neurons: such as RT, MRN and superior colliculus, motor related (SCm) in midbrain; (iv) from individuals to one hundreds of neurons, such as parabrachial nucleus (PB) and pontine reticular nucleus, caudal part (PRNc) in pons, gigantocellular reticular nucleus (GRN) and vestibular nuclei (VNC) in medulla, or to several hundreds of neurons, such as substantia nigra, compact part (SNc) in midbrain, IP and dentate nucleus (DN) in cerebellum.

**Fig. 2.**
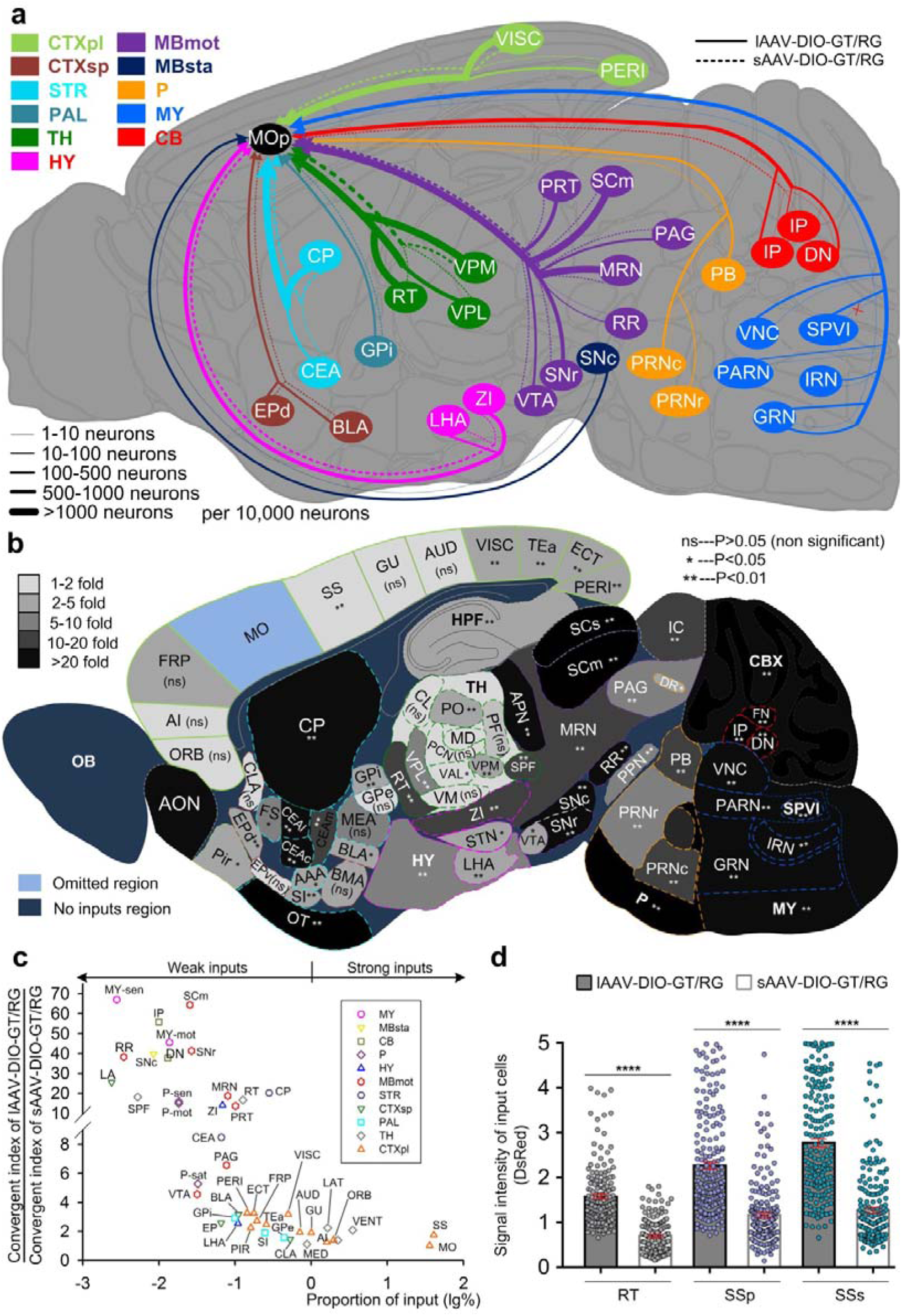
MAP-ENVIVDERS enables efficient tracing of input networks. **a,** Summary of representative sparsely traced input suberegions with current RV-mediated trans-monosynaptic tracing systems (sAAV-DIO-GT/RG, dashed lines), which were significantly enhanced by MAP-ENVIVIDERS (lAAV-DIO-GT/RG, solid lines). The thickness of each colored lines represented the average number of input neurons (n = 5 mice) in each subregions (per 10, 000 neurons). **b**, Summary of increased tracing efficiency in diverse brain regions with MAP-ENVIVDERS. n = 5 mice for each group. Mann-Whitney U test. n.s, non-significant (P > 0.05), *P < 0.05, **P < 0.01. **c**, Relationship between the proportion of inputs (input cells in each subregion relative to the total input cells across whole brain, which was denoted by logarithm to base 10 (lg %, x-axis) and the enhanced ratios provided by MAP-ENVIVIDERS (the ratio between convergent index of lAAV-DIO-GT/RG and convergent index of sAAV-DIO-GT/RG, y-axis). n = 5 mice for each group. **d**, Signal intensities of input neurons in reticular nucleus of the thalamus (RT), primary somatosensory area (SSp), and supplemental somatosensory area (SSs) labeled by lAAV-DIO-GT/RG-RV or sAAV-DIO-GT/RG-RV. n = 160 neurons for RT, n = 190 neurons for SSp and SSs, n = 3 mice for each group. two-tailed t-test ****P < 0.0001. All data are presented as mean ± s.e.m. Abbreviations: see **Supplementary Table 2**.

To further obtain specific enhanced ratios in diverse major regions and subregions achieved by MAP-ENVIVIDERS, the corresponding convergent index^8^ (the number of input cells divided by the number of starter cells) was calculated initially. Then, the ratio between convergent index of lAAV-DIO-GT/RG and convergent index of sAAV-DIO-GT/RG was plotted against the percentage of inputs (input cells in each subregion relative to the total input cells across whole brain). The relationship between the improvement and the labeling efficiency by MAP-ENVIVIDERS is clear: the poorer the labeling with the original system, the greater the improvement (**Fig. 2c**). We found that most of the originally underrepresented regions (< 1%) were enhanced by more than an order of magnitude and these regions were distributed throughout the brain, such as CP (20-fold) in striatum, RT (17-fold) in thalamus, ZI (14-fold) in hypothalamus. Brain regions with over 20-fold (20 to 60- fold) enhanced tracing efficiencies were more frequently found in midbrain, hindbrain and cerebellum, areas which were very far away from injection site, such as SNr (41- fold) and SNc (40-fold) in midbrain; reticular nucleus (gigantocellular: 46.0-fold and intermediate: 44-fold) in medulla; IP (56-fold) and DN (38-fold) in cerebellum, corresponding to the results of fluorescent images in **Fig. 1e**. Particularly, several brain regions in brain stem, including SCm, sensory related (MY-sen) and VNC in medulla, motor related (MY-mot), showed particular improvements of up to 70-fold (**Fig. 2b** and **Supplementary Fig. 4d**).

To examine whether brightness of input neurons could be enhanced by MAP-ENVIVIDERS, we further compared the signal intensities of the input neurons traced by lAAV-DIO-GT/RG and sAAV-DIO-GT/RG in the following brain regions: SSp, supplemental somatosensory area (SSs) and RT (**Supplementary Fig. 1a**, right panel). We found that the signal intensities of the input neurons were significantly enhanced in lAAV-DIO-GT/RG labeling groups compared with sAAV-DIO-GT/RG labeling groups (**Fig. 2d** and **Supplementary Fig. 4e**; 2.27-fold for RT, 1.94-fold for SSp and 2.19-fold for SSs; unpaired t-test, P < 0.0001, n = 3 mice). These data collectively showed that MAP-ENVIVIDERS significantly enhanced not only the number but also the brightness of the labeled input neurons. Taken the facts of similar number of starter cells, significantly enhanced number and fluorescent intensity of input cells suggested that many more RVs are produced by the starter cells in MAP-ENVIVIDERS.

### Demonstration of MAP-ENVIVIDERS for reliable, density-controllable super-bright neuronal labeling

Dissection of neurocircuits on individual neuron level is critical for understanding neuronal structure-function relationship and cell-type classification^11^. In previous studies, dual rAAV-based Cre-lox/Flp-FRT recombinase systems and the tTA/TRE transactivation system have been successfully developed to label limited or tunable numbers of neurons of specific cell types^13, 20, 21^. However, a viral copackaging strategy has never been explored in these systems. Therefore, we next explored whether MAP-ENVIVIDERS could improve the labeling brightness for mapping projectomes of individual neurons. We first validated MAP-ENVIVIDERS in wild-type mice with copackaging a Cre-expressing vector and a Cre-dependent vector (**Fig. 3a**). We generated three littermate rAAVs (abbreviated for lAAVs hereafter) with a CMV promoter-mediated Cre-expressing vector (AAV-CMV-Cre) and a Cre-inducible, double-floxed EYFP element (AAV-double floxed-EYFP)^34^ at ratios of 1:20,000 (lAAV^20,000^), 1:200,000 (lAAV^200,000^) and 1:1,000,000 (lAAV^1,000,000^). We found that these three lAAVs produced different labeling densities, but with similar brightness in the soma and long-range axonal projections when injected into MOp (**Fig. 3b** and **Supplementary Fig. 6**). The labeled somas were well separated, and the fine structures, including various types of spines^35^ and boutons on local and long-range projecting arborizations^15^, could be clearly visualized and identified (**Supplementary Fig. 5**).

**Fig. 3.**
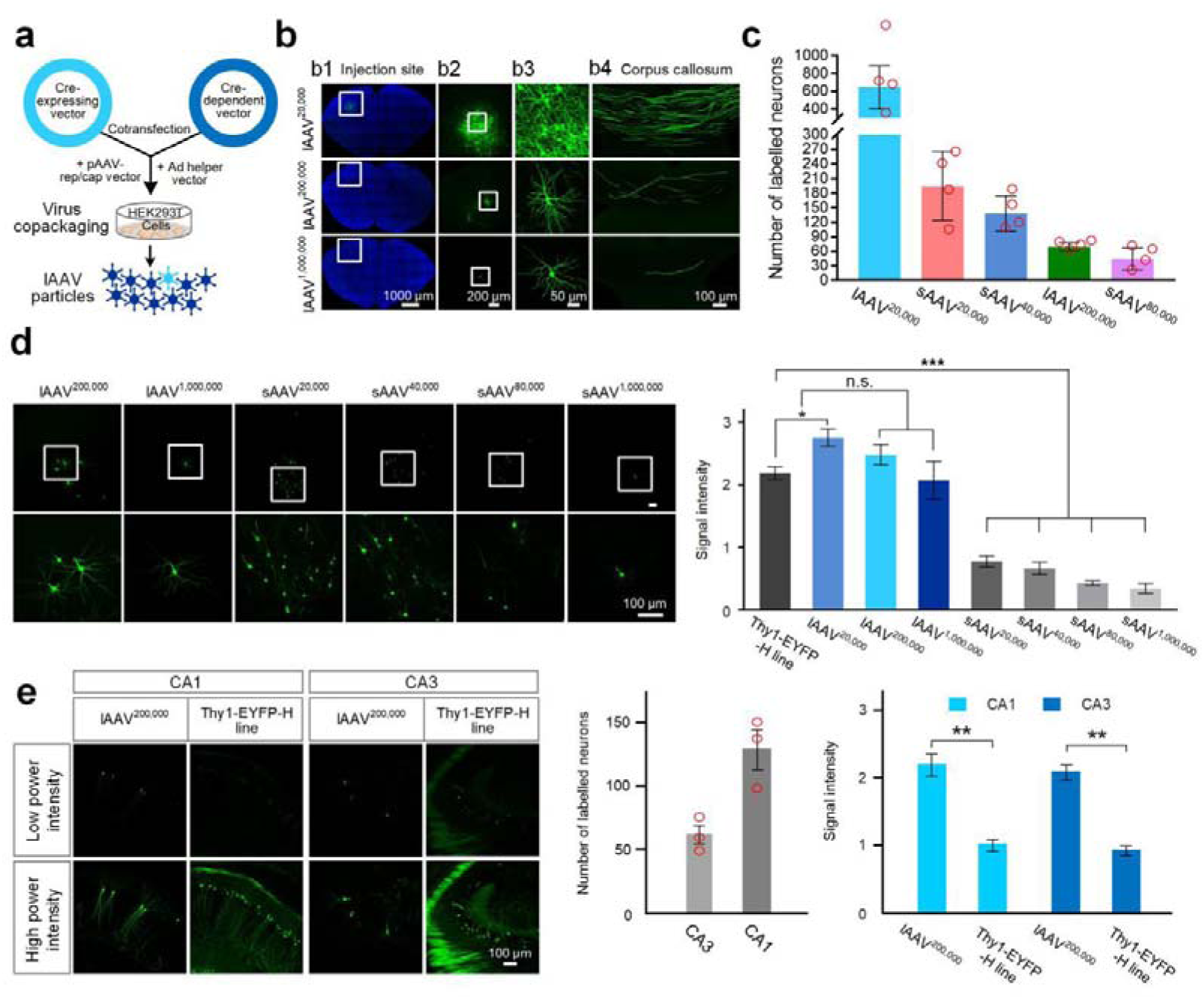
MAP-ENVIVIDERS enables density-controllable, super-bright neuron labeling in wild-type mice. **a**, Schematic for the production of lAAVs. Briefly, a Cre-expressing vector and a Cre-expressing vector were mixed at preset ratios on cellular level, generating lAAV particles. **b**, Representative confocal images of neurons from mice injected with 100 nl of lAAVs produced by copackaging pAAV-CMV-Cre and pAAV-EF1α-double floxed-EYFP at the indicated ratios. Images show neurons at the injection sites (**b1-b3**) and their long axonal projections in the corpus callosum (**b4**). **c**, Quantification of neurons labeled by different lAAVs and sAAVs (n = 4 mice each). **d,** Comparison of brightness between neurons labeled by different lAAVs and sAAVs. Upper panel: confocal images; lower panel: quantifications of signal intensities of the labeled somas. Neurons from the Thy1-EYFP-H mouse line were used as a reference (n = 40 cells for the Thy1-EYFP-H mouse line; n = 40 cells for lAAV^20,000^, lAAV^200,000^, sAAV^20,000^, sAAV^40,000^, and sAAV^80,000^ from 3 mice; n = 20 cells for lAAV^1,000,000^ and n = 17 cells for sAAV^1,000,000^, from 4 mice). One-way ANOVA with Dunnett’s post hoc test. **e**, Applications of lAAV^200,000^ in CA1 and CA3. Comparisons of brightness of lAAV^200,000^-labeled neurons in CA1 and CA3 (n = 40 cells for each from 3 mice) with Thy1-EYFP-H mouse line (n = 40 cells) were shown with confocal images (left panel) and measurement of signal intensities (right panel) (n = 40 cells for each from 3 mice). Two-tailed t-test. Middle panel, quantifications of labeled neurons in CA1 and CA3 (n = 3 mice each). All data are presented as the mean ± s.e.m. n.s, non-significant (P > 0.05), *P < 0.05, **P < 0.01, ***P < 0.001. Abbreviations: see **Supplementary Table 2.**

To compare the differences in labeling efficiency and brightness between lAAVs and mixtures of independently packaged rAAVs (sAAVs), we performed parallel experiments with mixtures of Cre-expressing rAAV and Cre-dependent rAAVs at ratios of 1:20,000, 1:40,000, 1:80,000 and 1:1,000,000 based on previous methods^12^ (abbreviated for sAAV^20,000^, sAAV^40,000^, sAAV^80,000^ and sAAV^1,000,000^, see Methods). We selected Thy1-EYFP-H mice^36^, one of widely used transgenic lines as reference. Although density-controllable labeling could be achieved in both lAAVs and sAAVs- based systems, labeling by lAAVs was significantly brighter and more efficient under the same experimental conditions (**Fig. 3c** and **d**, left panel). Specifically, the brightness of sparsely labeled neurons by lAAVs (**Supplementary Fig. 1c,d**) was equivalent to Thy1-EYFP-H mice, but ∼3 times stronger for lAAV^200,000^ compared with sAAV^20,000^ and ∼7 times for lAAV^1,000,000^ compared with sAAV^1,000,000^, respectively (P < 0.001), demonstrating the sensitivity of MAP-ENVIVIDERS (**Fig. 3d**, right panel and **Supplementary Fig. 7**). In accordance with previous study^31^, the Cre titers of lAAVs and sAVVs were similar as measured by quantitative polymerase chain reaction, however, the number of labeled neurons for lAAV^20,000^ was nearly 4- fold that of sAAV^20,000^, demonstrating the efficiency of MAP-ENVIVIDERS (**Fig. 3c** and **Supplementary Table 3**). Similar results were also obtained in other brain regions, such as CA1 and CA3, where the labeled neurons were significantly brighter than those in the Thy1-EYFP-H mouse line (**Fig. 3e**, left and right panel; **Supplementary Fig. 8**). However, the number of neurons labeled differed among the brain regions for the same lAAV preparation (**Fig. 3e**, middle panel). These results collectively showed that the MAP-ENVIVIDERS labeling strategy is significantly more sensitive, efficient and reproducible than methods using mixtures of independently packaged rAAVs for density-controllable neuron labeling.

### MAP-ENVIVIDERS enables identification of novel type of neurons

To further explore the potential of MAP-ENVIVIDERS in obtaining brain-wide long-range projections, we used fluorescence micro-optical sectioning tomography (TDI-fMOST)^37^ to automatically image the brain sample injected with lAAV^200,000^ in the MOp of wild-type mouse. Imaging was conducted with a voxel size of 0.176 × 0.176 × 1 µm^3^, generating 12,841 continuous coronal section across the whole brain (**Fig. 4a** and **Supplementary Fig. 9**). Among the 11 reconstructed MOp pyramidal neurons, four intratelencephalic neurons located in L2, L3, and L6a projected to the ipsi- and contralateral striatum and cortex, and four pyramidal tract neurons located in L5b projected to diverse subcortical brain regions, as previously reported^38–42^ (**Fig. 4c**; **Supplementary Fig. 10a,b** and **Supplementary Video 1**). Interestingly, the remaining three neurons (No. **s1**-**s3**) formed a novel type of pyramidal neuron located in L5b (**Fig. 4b** and **Supplementary Fig. 10c**). We termed these “boomerang neurons” because they send a collateral axonal branch to the contralateral MOp and dorsal caudoputamen (CP) via the corpus callosum, which then returns to the ipsilateral side via the anterior commissure, innervating the ipsilateral CP region via a collateral on the ipsilateral side, thus forming a symmetrical track (**Fig. 4d,e**; **Supplementary Fig. 10d** and **Supplementary Video 2-4**). All of these reconstructed boomerang neurons had complex axonal projections, with over axonal 1,200 branches totaled at ∼22 cm long on average (**Supplementary Fig. 14**). Therefore, the use of MAP-ENVIVIDERS together with TDI-fMOST allowed us to document the projectome of individual neurons in great detail as well as to identify novel type of neurons.

**Fig. 4.**
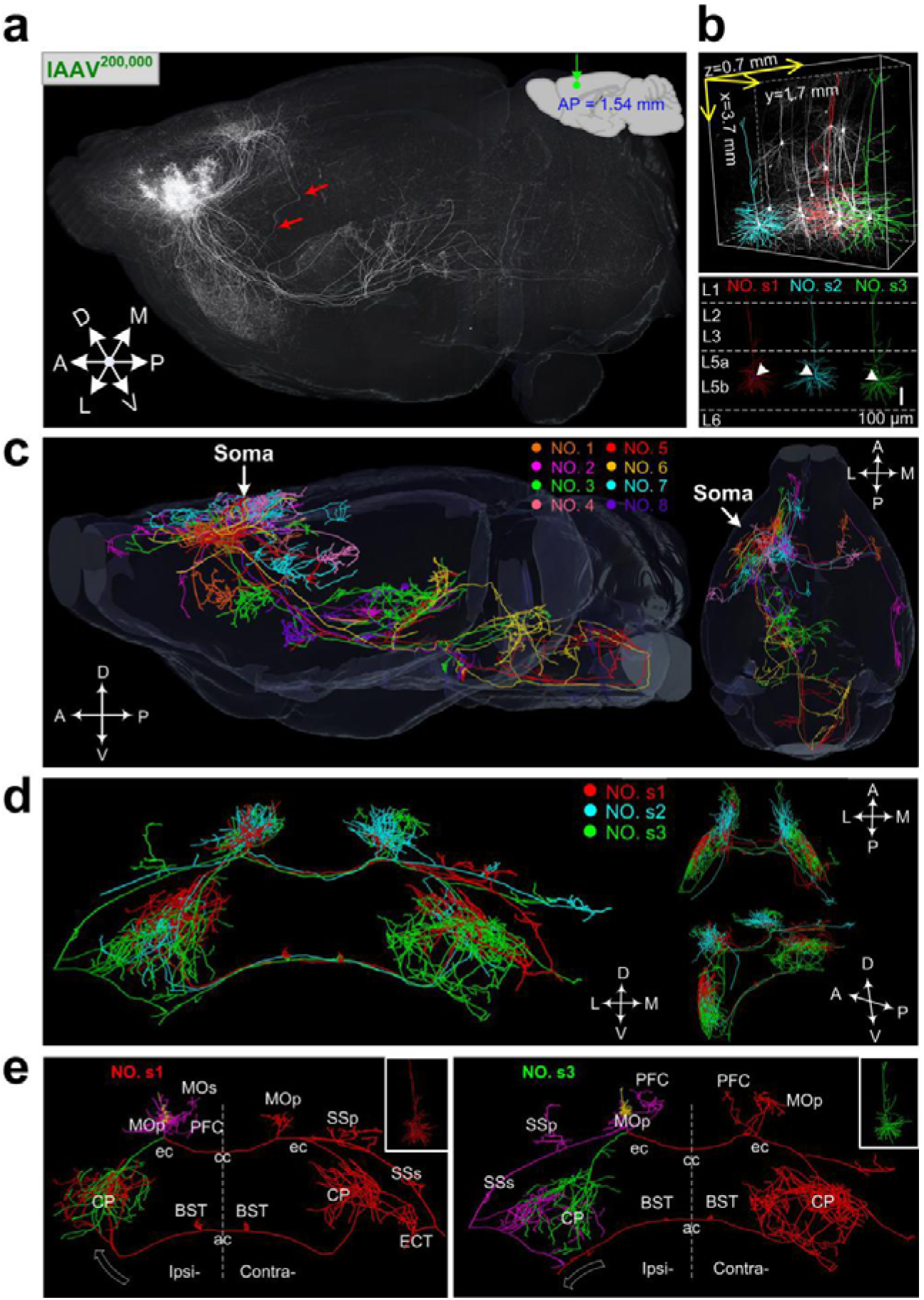
MAP-ENVIVIDERS enables identification of a novel type of neuron in MOp. **a**, Sagittal view showing lAAV^200,000^-labeled MOp whole-brain dataset. The red arrows indicate axon pathway of boomerang neurons. **b**, Raw signals (top) and reconstructed dendritic morphologies (bottom) of three boomerang neurons (No. **s1-s3**). **c**, Summary of axonal morphologies of the eight reconstructed pyramidal neurons (**No. 1-8**) in sagittal (left panel) and horizontal (right panel) views. **d**, Coronal (left), horizontal (upper right), and sagittal (with a rotated angle, lower right) views showing axonal morphologies of three boomerang neurons. **e**, Localization of the brain-wide axonal projections of neurons **s1** and **s3** (dendrites: yellow; local axons: magenta; the left and right branches originating from the ipsilateral external capsule are shown in green and red, respectively). Dashed white arrows indicate the termination directions of the main axons. Abbreviations: see **Supplementary Table 2.**

### MAP-ENVIVIDERS enables unbiased, cell-type-specific, sparse and super-bright labeling among diverse transgenic mice

Cell-type-specific labeling is crucial for dissecting the functional role and circuitry of a given type of cell^21^. To demonstrate the potential of MAP-ENVIVIDERS for efficient cell-type-specific and density-controllable labeling with multiple Cre-driver lines, we next redesigned MAP-ENVIVIDERS with an Flp-expressing vector (e.g., AAV-EF1α-Flp) and an both Flp- and Cre-dependent vector (e.g., AAV-hSyn Con/Fon EYFP^43^) and copackaged two vectors at different ratios to generate cslAAV^2,000^ and cslAAV^20,000^ (cs refers to cell-type-specific hereafter, **Fig. 5a**). For validations, we injected the resulted two cslAAVs into MOp of Thy1-Cre transgenic mice and found that tunable neuron labeling was achieved with different ratios (**Supplementary Fig. 11a,b**), indicating the functionality of MAP-ENVIVIDERS with Cre-driver transgenic lines.

**Fig. 5.**
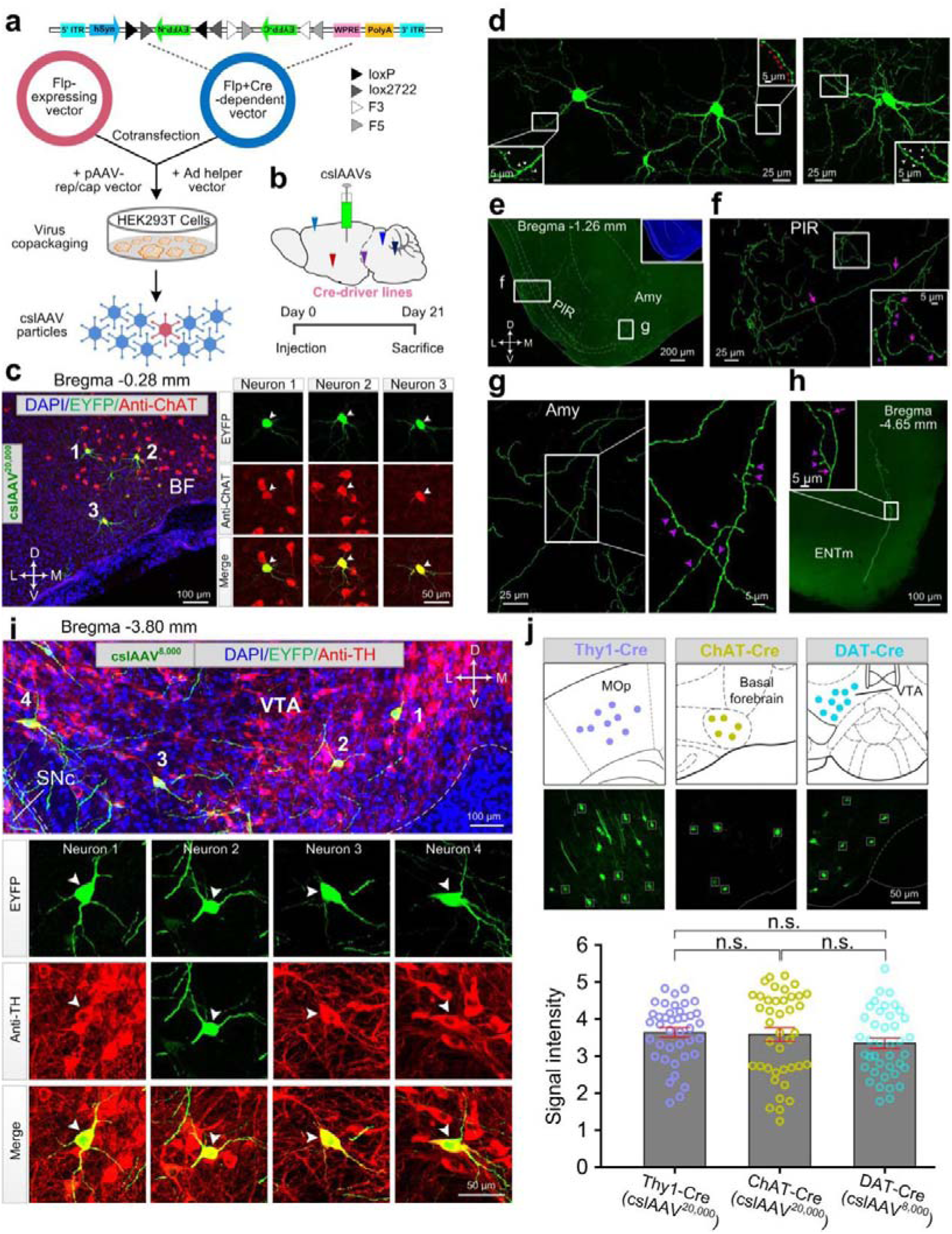
MAP-ENVIVIDERS enables unbiased, cell-type-specific, sparse and super-bright labeling among diverse transgenic mice. **a**, Schematic showing the generation of cslAAVs and the Cre- and Flp-dependent AAV vector. ITR, inverted terminal repeat; hSyn, human synapsin promoter; WPRE, woodchuck hepatitis virus post transcriptional regulatory element. **b**, Schematic of the experiment for the data in **c**-**i**. **c**, Low-magnification (left) and high-magnification (right) images showing colocalization of three cssAAV^20,000^-labeled basal forebrain cholinergic neurons (neurons 1 to 3, green) with anti-choline acetyltransferase (ChAT) antibody staining (red). **d**, Fine structure of three cholinergic neurons in **c**. Insets are enlargements of the corresponding boxes, white arrowheads indicate spines, and red arrowheads indicate boutons. **e-h**, Representative images showing long-range axonal projections in the piriform area (PIR, **f**), amygdala (Amy, **g**), and entorhinal area, medial part, dorsal zone (ENTm, **h**). **i**, low-magnification (upper panel) and high-magnification (lower panel) images showing immunostaing of four cssAAV^8,000^-labeled dopaminergic neurons (neurons 1 to 4, green) in ventral tegmental area (VTA). Anti-tyrosine hydroxylase (TH) antibody staining, the definitive marker for dopaminergic neurons, were shown in red. **j**, pper panel, schematic for measurement (upper panel) and comparisons (lower panel) of signal intensities of individual neurons labeled by different cslAAVs in multiple Cre-driver transgenic lines, including cssAAV^20,000^-labeled Thy1-Cre mice, cssAAV^20,000^-labeled ChAT-Cre mice and cssAAV^8,000^-labeled DAT-Cre mice. (n = 40 cells from 3 mice for each). One-way ANOVA with Turkey’s post hoc test. All data are presented as the mean ± s.e.m. n.s, non-significant (P > 0.05). Abbreviations: see **Supplementary Table 2.**

To achieve cell-type-specific sparse and super-bright labeling of cholinergic neurons, we next applied one of cslAAVs, cslAAV^20,000^ into the basal forebrain (BF) of choline acetyltransferase (ChAT)-Cre transgenic mice^23^ (**Fig. 5b**). Among 54 ± 4 well-separated neurons labeled with cslAAV^20,000^ (n = 4 mice), 97.6 ± 1.6% of which were ChAT positive (**Fig. 5c**, **Supplementary Fig. 12** and **Supplementary Table 3**). Fine structures of these sparsely neurons, both near and far from the somas, could be visualized clearly (**Fig. 5d-h**) and their fine long-range projections covered diverse brain regions as previously reported^44^ (**Supplementary Fig. 13**). Parallel experiments with wild-type mice did not label any cells (n = 4 mice; **Supplementary Fig. 11c**), confirming the cell-type specificity of the method. In another set of parallel experiments in which cssAAV^2,000^ and cssAAV^20,000^, mixtures of the corresponding independently packaged rAAVs (see Methods) were injected into BF of ChAT-Cre transgenic mice. We found that no neurons were labeled with cssAAV^20,000^ and only a few faintly labeled with cssAAV^2,000^ (n = 5 mice each; **Supplementary Fig. 11d,e**). These results demonstrated that the MAP-ENVIVIDERS strategy was significantly more sensitive and efficient for cell-type-specific neuron labeling.

The strength and abundance of Cre recombinase varied greatly among diverse Cre-driver transgenic lines^22, 23^. To examine the labeling efficiencies in different Cre-driver mice, we measured the fluorescent intensity of individual neurons labeled by cslAAVs with different labeling densities in three Cre-driver lines under the control of different promoters, including Thy1 promoter (Thy1-Cre), dopamine transporter promoter (DAT-Cre)^45^ and choline acetyltransferase promoter (ChAT-Cre) (**Supplementary Fig. 1c** and **Supplementary Table 3**). We found that averaged fluorescent intensity of individual neurons in showed no apparent difference among these three different types of Cre-driver transgenic mice (**Fig. 5i,j**). These results showed that the MAP-ENVIVIDERS labeling strategy enables unbiased, cell-type-specific, sparse and super-bright labeling among diverse transgenic mice.

### MAP-ENVIVIDERS enables identifying presynapses of individual cell-type-specific neurons

To reveal fine morphologies of individual cholinergic neurons, we also performed whole-brain mapping of BF-injected brains of ChAT-Cre mice (**Fig. 6a**). Based on cell-type-specific, sparse but super-bright labeling, four cholinergic neurons we reconstructed showed distinctive projectomes but all shared with extremely complex axonal aborizations, with an average of 4,059 axonal branches (**Fig. 6b; Supplementary Fig. 14** and **Supplementary Video 5**). Remarkably, we obtained an individual cholinergic neuron (No. **c2)** containing incredible ∼9,500 axonal branches, which were nearly 9-times more than previously reconstructed individual cholinergic neuron by the combination of drug and transgenic mice strategy^46^. Particularly, this neuron contained two major axonal branches with one of them send massive arborizations dominating layer 2 to layer 4 of the ipsilateral anterolateral/lateral visual area (VISal/l, ∼7,840 axonal branches), a critical region for visual processing^47^, and the other terminated in ipsilateral subiculum (SUB) (**Fig. 6e,f** and **Supplementary Video 6**). Projections of cholinergic neurons to hippocampus have long been known as a key player in the formation of learning and memory^48^. To this end, we reconstructed two BF cholinergic neurons (No. **c3** and No. **c4**) which send projections covering entire hippocampus (−1.0 to −3.8 mm from the bregma; **Fig. 6c**). Specifically, the former (No. **c3**) exhibited a point to point projection mode with sole projections to contralateral dorsal and ventral hippocampal region (HIP) from BF, whereas the latter (No. **c4**) projected to several cortical regions successively, including ipsilateral anterior piriform area, prefrontal cortex, motor areas and somatosensory areas and finally terminated in ipsilateral hippocampus with massive massive arborizations in ventral HIP followed by less axonal branches in dorsal HIP and a single axonal branch regulating both ipsilateral and contralateral dorsal HIP (**Fig. 6b,c**). Similar to No. **c3**, the fourth neuron (No. **c4**) also exhibited a point to point projection mode with projections from BF to ipsilateral CP (BF→CP) in a fanwise manner (−1.1 to −2.1 mm from the bregma; **Fig. 6d**). These results demonstrate that the MAP-ENVIVIDERS could capture morphological details of individual neurons with cell-type specificity to a great extent.

**Fig. 6.**
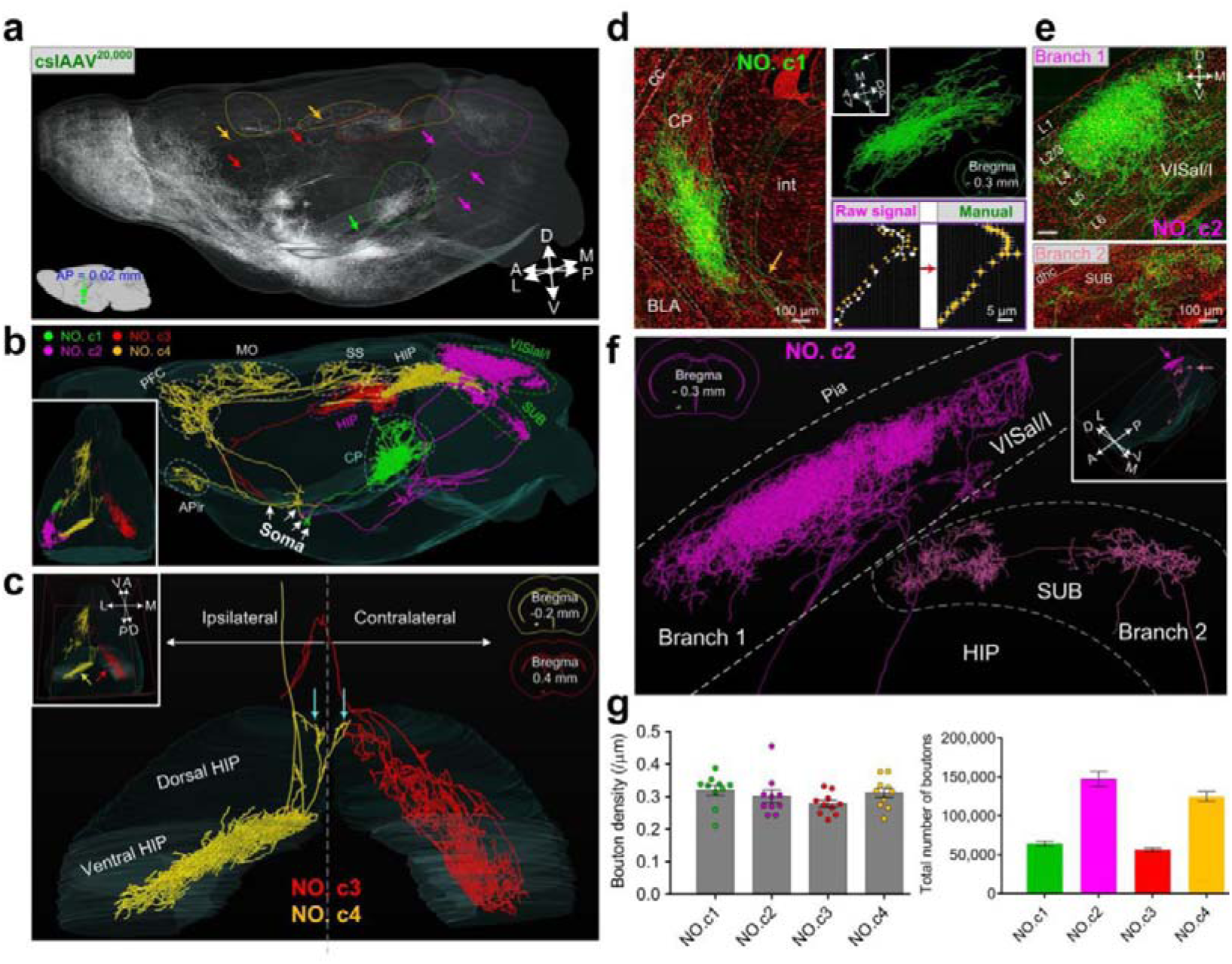
MAP-ENVIVIDERS enables identification of presynapses of individual cell-type-specific neurons. **a**, Sagittal view showing whole-brain labeling after injection of cslAAV^20,000^ into the basal forebrain. Arrows indicate the main axons and colored circles indicate the terminal arborizations of each reconstructed neuron. **j**, Summary of the axonal morphologies of four reconstructed cholinergic neurons (**No. c1-c4**) in sagittal and the horizontal view (insets). **c**, Three dimensional view showing terminal arborizations (indicated by arrows in upper left insets) in dorsal and ventral hippocampal region (HIP) of two cholinergic neurons (**No. c3**: red; **No. c4**: yellow). Cyan arrows indicated a single axonal branch of **No. c4** with concomitant regulation of both ipsilateral and contralateral dorsal HIP. **d**, Left panel, raw signal with propidium iodide (PI)-stained cytoarchitecture reference of neurons c1 (green: maximum intensity projection of 1,000 coronal sections; Red: 5 µm thickness). Yellow arrow indicated the main axon. Upper right panel: reconstructions of complex axonal arborizations in CP. Lower right panel: enlargements of box region in upper left panel demonstrating schematic for manual bouton counting in **g**. **e**, Raw signal with PI-stained cytoarchitecture reference showing complex axonal arborizations in the anterolateral/lateral visual area (VISal/l; branch 1; upper) and subiculum (SUB; branch 2; lower) of neurons **c2**. Branch 1 and branch 2 were maximum intensity projection of 1,800 and 400 coronal sections (green signal), respectively. Red signals were 5 µm thickness for both. **f**, Reconstructions of branch 1 and branch 2 in **e**. Brain outlines in **c**,**d**,**f** depicted approximate somal locations of respective neurons. **g**, Averaged bouton density (left panel, n = 10 ROIs for each neuron) and estimated total number of boutons (right panel) of four reconstructed cholinergic neurons. Each circle represents one ROI. Abbreviations: see Supplementary Table 2.

Plasticity of presynaptic axonal boutons served as key elements for the maintenance of neural circuit functions and loss of bouton density has been associated with several neurodegenerative disorders^49, 50^. To assess the total number of individual cholinergic neurons, we randomly selected ten regions of interest (ROIs) from raw signal of each cholinergic neuron and counted the boutons along the axon shafts manually (lower right panel in **Fig. 6d** and **Supplementary Fig. 15**; also see methods). The averaged bouton density (defined as bouton numbers over axonal length) of four cholinergic neurons was estimated to be 0.303 ± 0.016 bouton/μm (No. **c1**: 0.320 ± 0.016 bouton/μm; No. **c2**: 0.302 ± 0.020 bouton/μm; No. **c3**: 0.279 ± 0.011 bouton/μm; No. **c4**: 0.312 ± 0.015 bouton/μm; n = 10 ROIs for each neuron; left panel in **Fig. 6g** and **Supplementary Table 4**). Based on these results, we thus estimated that averaged total number of four cholinergic neurons was 98,496 ± 5,230 (No. **c1**: 64,295 ± 3,174; No. **c2**: 147,858 ± 9,668; No. **c3**: 56,592 ± 2,151; No. **c4**: 125,138 ± 5,928; right panel in **Fig. 6g** and **Supplementary Table 3**), which (to the best of our knowledge) has not been reported previously. Such huge number of boutons obtained by MAP-ENVIVIDERS indicated that an individual cholinergic neuron enjoyed a wide range of modulation coverage.

### Copackaging of rAAVs significantly reduces their mutual suppression

Since we demonstrated that MAP-ENVIVIDERS significantly improved the tracing efficiencies of input networks as well as enhanced the labeling brightness of output networks, we next made initial attempts to investigate probable mechanisms for the powerfulness of this viral copackaging-based labeling strategy. Previous studies have indicated that the interactions among different viral vectors, such as AAV over herpes simplex virus^24^, AAV over adenovirus^25^, or even among the same types of viruses (e.g., herpes simplex virus and pseudorabies virus)^26–28^, can severely affect their gene expressions. We next evaluated the degree of interaction between two different rAAVs. To eliminate the effects of other factors, we used two independently packaged, highly similar rAAVs, which were same serotyped (rAAV2/9) under the control of the same ubiquitous promoter (EF1α) but only expressing different fluorescent proteins (EYFP or mCherry). We injected equal ratio mixture of two rAAVs (abbreviated for sAAV-EYFP/mCherry for convenience, s refers to stranger hereafter) into the MOp of wild-type mice. For the controls, a parallel procedure was followed but with rAAV2/9-EYFP or rAAV2/9-mCherry mixed equally with phosphate buffered saline (PBS). The fluorescence intensity was significantly lower for both EYFP and mCherry when the vectors were used in combination than when used separately (**Fig. 7a,b**), by a factor of 2.4 for EYFP and 1.4 for mCherry (**Fig. 7c**), demonstrating significant mutual suppression among different rAAVs.

**Fig. 7.**
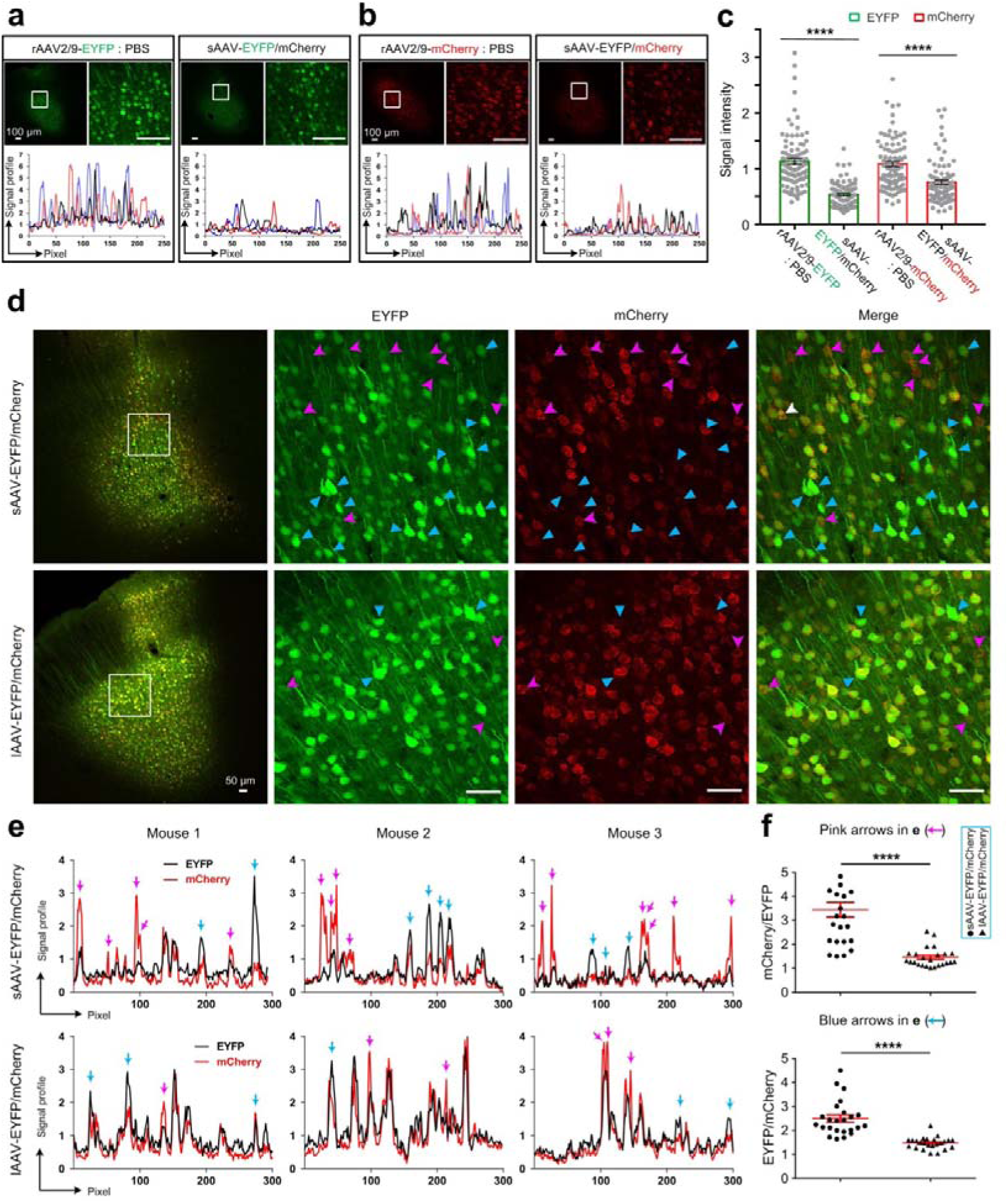
Viral copackaging reduces rAAV suppression. **a-b**, Representative confocal images and signal profiles of EYFP and mCherry in neurons of the MOp of wild-type mice injected with 100nl 1:1 mixtures of rAAV2/9-EYFP : PBS, rAAV2/9-mCherry : PBS and rAAV2/9-EYFP : rAAV2/9- mCherry (sAAV-EFYP/mCherry). **c**, Fluorescence intensity quantified in 90 neurons (n = 3 animals) from **a** and **b**. **d**, Representative confocal images showing expression of EYFP and mCherry in neurons of MOp of wild-type mice injected with 100nl sAAV-EFYP/mCherry (upper panel) or lAAV- EFYP/mCherry (lower panel). Pink and cyan arrowheads indicate neurons with higher EYFP and mCherry signals, respectively. **e**, Representative signal profiles of mCherry and EYFP in three mice injected with sAAV-EFYP/mCherry (upper panel) or lAAV-EFYP/mCherry (lower panel). Pink and cyan arrows indicate neurons with higher EYFP and mCherry signals, respectively. **f**, Ratio of mCherry/EYFP (pink arrows) and EYFP/mCherry (cyan arrows) fluorescence intensity in **e**. All data are presented as the mean ± s.e.m., n = 3 animals for each group in **a**-**f**. Two-tailed t-test, ****P < 0.0001.

Previous studies have employed viral copackaging strategy for cost- and time- effective AAV manufacturing^29, 30^. We next examined whether viral copackaging strategy could increase homogeneity over independently packaged rAAVs. To achieve this, we copackaged the two corresponding plasmids at a 1:1 ratio and then injected the resulting rAAVs (abbreviated as lAAV-EYFP/mCherry, where l refers to littermate hereafter) into MOp as described above. Equivalent expressions of EYFP and mCherry within individual cells were observed more frequently in neurons co-infected by lAAV-EYFP/mCherry than in neurons coinfected by sAAV-EYFP/mCherry, which instead tended to produce unbalanced expression of EYFP and mCherry (**Fig. 7d,e**, top vs. bottom rows). For neurons with unbalanced expressions (arrows in **Fig. 7e**), The ratio of mCherry/EYFP and EYFP/mCherry fluorescence intensity with higher mCherry and EYFP signals, respectively, was significantly higher in neurons injected with sAAV-EYFP/mCherry than in those injected with lAAV-EYFP/mCherry (mCherry/EYFP ratio (pink arrows): 3.44 ± 0.76 vs. 1.46 ± 0.20, respectively, and EYFP/mCherry ratio (cyan arrows): 2.50 ± 0.37 vs. 1.48 ± 0.12, respectively; mean ± s.e.m, unpaired t-test, P < 0.0001, n = 3 mice, **Fig. 7f**). These results indicated that copackaging rAAVs can substantially reduce mutual suppression and significantly enhance compatibility among different rAAVs.

## DISCUSSION

In the present study, we developed MAP-ENVIVIDERS and generated series of vector mixing-based rAAV cocktail providing more efficient cell-type-specific whole-brain input mapping and improved super-bright and sparse labeling for certain types of neurons. We then found that strong mutual suppression existed among independently packaged rAAVs and copackaging of rAAV-producing plasmids enhanced co-expression of genes among multiple rAAVs.

Based on our results, we concluded that MAP-ENVIVIDERS were more prominent in complicated systems requiring co-expression of multiple rAAVs compared with previous strategy achieved by co-injections or co-administrations of several seperately packaged rAAVs. For non-cell-type-specific super-bright and sparse labeling (co-expression of Cre and Cre-dependent fluorescent protein), MAP-ENVIVIDERS provided ∼3-fold more labeled neurons and ∼5-fold more bright fluorescent intensity (**Fig. 3c,d**). For cell-type-specific super-bright and sparse labeling (co-expression of Flp, Cre and Flp-/Cre-dependent fluorescent protein), MAP-ENVIVIDERS provided more than 10-fold increase of labeling efficiency (**Fig. 5c** and **Supplementary Fig. 11d**,**e**). Thus, MAP-ENVIVIDERS could achieve the same efficiency with much lower dose of virus, which may reduce cytotoxicity and experimental cost. Notably, since MAP-ENVIVIDERS is very sensitive, careful consideration of mixing ratios for viral copackaging was needed to label desired number of neurons. For cell-type-specific super-bright/sparse labeling, the strength and abundance of Cre recombinase varies greatly among different cell types and brain regions^22, 23^, thus the mixing ratio of Flp and Flp-/Cre-dependent fluorescent protein need to be carefully tested.

Moreover, when MAP-ENVIVIDERS was applied for circuit tracing which involves not only compatibility two different rAAVs but coherence between two labeling systems (i.e., rAAVs-RV combination), labeling efficiency was dramatically increased (10- to 70-fold) for 40% of the input brain regions, which were sparsely labeled by previous strategies (**Fig. 1,2**). The much stronger DsRed signals and the larger number of total input neurons suggested that more RVs may be produced with the MAP-ENVIVIDERS strategy. This is probably due to the reduction of mutual suppression between the rAAVs expressing TVA and RG. The AAVs in the starter cells expressed more TVA and RG, leading to more entry and assemble of infectious RVs and eventually higher transsynaptic tracing efficiency (convergent index from averaged 32 to 52, **Supplementary Table 1**). Moreover, it is feasible to combine MAP-ENVIVIDERS with optimized RG (oG)^10^ and other RV strains (e.g., the CVS-N2c^ΔG^ strain)^9^ to further improve the tracing efficiency. Further efforts towards these largely enhanced input neurons in diverse brain regions labeled by MAP-ENVIVIDERS including identifications the components of cell types with the aid of immunochemistry (homogeneous to the cell types labeled with current RV-mediated trans-monosynaptic tracing systems or not) and electrophysiology- or optogenetics-assisted functional studies.

Among diverse Cre-driver transgenic line for cell-type-specific neuronal labeling, Thy1 promoters-mediated lines exhibited the most robust gene expression levels^37^. With MAP-ENVIVIDERS, the fluorescent intensity of sparsely labeled neurons in choline acetyltransferase promoter- and dopamine transporter promoter-mediated lines approaching that achieved by Thy1 promoters (**Fig. 5j**). This unbiased property for cell-type-specific, sparse and super-bright labeling highlights the great sensitivity of MAP-ENVIVIDERS strategy. Combined with whole-brain imaging systems, MAP-ENVIVIDERS demonstrated great advantages for capturing more details of individual cell-type-specific neurons (e.g. an individual cholinergic neuron with ∼9,500+ axonal branches and averaged 148,000 presynaptic axonal boutons) (**Fig. 6**). To obtain exact number of boutons of specific individual neurons faithfully, further efforts are needed including co-expressing fluorescent proteins with bouton indicative marker, such as well-known synapophysin^51^ along the axons, as well as developing automatic or semi-automatic softwares for bouton identification^52^.

Though further efforts are needed to explore the deeper mechanisms leading to the high sensitivity and efficiency of MAP-ENVIVIDERS, our initial attempts indicated that strong mutual suppression existed among different rAAVs, a phenomenon similar to other kinds of viruses, such as herpes simplex virus, hepatitis C virus and influenza A virus^53, 54^ (**Fig. 7a-c**). We further found that viral copackaging strategy significantly ameliorated this suppression and enhanced compatibility of multi-gene expressions in the same cells (**Fig. 7d-f**), the probable reasons for which were explained as follows: (i) Previous studies have demonstrated that the purities of rAAVs was a key determinant for their infection efficiency^55–57^. Viral copackaging strategy will reduce the heterogeneity of impurities, including cyto-toxins, cellular fragments and proteins, and culture media residues deprived from viral preparations of independently packaged rAAVs, thus reduce effect of strong immune responses on the state of the host cells near the injection site. (ii) Gene cassettes delivered by cocktails of rAAVs with different serotypes may elicit different immune responses, leading to varying reductions in gene expression^58, 59^, which could also be avoided by viral copackaging. Thus, the amplification of signals by recombinase systems combined with effects of (i) and (ii) reasonably leads to significant improvements of efficiency.

Besides the anatomical studies, MAP-ENVIVIDERS is suitable for functional studies of neural circuitry^60, 61^ and may even be used in other organs beyond mammal brain. Furthermore, MAP-ENVIVIDERS also provides great potential to facilitate the co-expression of multiple genes and reduce the harmful effects for clinical gene therapy^61^.

## METHODS

### Animals

All animal experimental procedures were approved by the Institutional Animal Ethics Committee of Huazhong University of Science and Technology (HUST) and Wuhan Institute of Physics and Mathematics (WIPM). All mice used in this study were 2-month-old male mice, including C57BL/6J mice, Thy1-EYFP-H line transgenic mice (Jackson Laboratory, stock number 003782, USA), Thy1-Cre transgenic mice (Jackson Laboratory, stock number 006143, USA), ChAT-Cre transgenic mice (Jackson Laboratory, stock number 006410, USA) and DAT-Cre transgenic mice (Jackson Laboratory, stock number 006660, USA).

### Virus preparations

All viruses used in this study were customized or commercially provided by BrainVTA Science and Technology Company (Wuhan, China). Three general types of viruses were used: copackaged littermate AAVs (lAAVs, l refers to littermate) for enhancing viral compatibility and coherence among different viruses, independently packaged stranger AAVs (sAAVs, s refers to stranger) as control, and genetically modified RV (EnvA-SADΔG-DsRed) for retrograde trans-monosynaptic labeling. Production of rAAVs and genetically modified RV were prepared as previously described^63, 64^.

Both general types of AAVs were serotype 2/9. They were used for four purposes: (i) non-cell-type-specific; (iii) cell-type-specific neuronal labeling; (iii) trans-monosynaptic labeling of the direct input network of specific type of starter cells as well as (v) verification of viral interactions, enhancing infection and exogenous gene expression efficiency.

For purpose (i), the plasmids were AAV-EF1α-double floxed-EYFP-WPRE-HGHpA^34^ (Addgene #20296, a generous gift from Dr. Karl Deisseroth, Stanford University) and AAV-CMV-Cre. The littermate rAAVs were lAAV^20,000^, lAAV^200,000^ and lAAV^1,000,000^. As controls, one of the two stranger rAAVs, rAAV2/9-CMV-Cre, was initially diluted to ratios of 1:20,000, 1:40,000, 1:80,000 and 1:1,000,000 in phosphate-buffered saline (PBS) before mixed equally with the other stranger rAAV, rAAV2/9-EF1α-double floxed-EYFP based on previous methods^12^ (the resulting rAAVs were abbreviated for sAAV^20,000^, sAAV^40,000^, sAAV^80,000^ and sAAV^1,000,000^, respectively).

For purpose (ii), the plasmids were AAV-hSyn Con/Fon EYFP^43^ (Addgene #55650, a generous gift from Dr. Karl Deisseroth, Stanford University) and AAV-EF1α-Flp. The littermate rAAVs were cslAAV^2,000^, cslAAV^8,000^ and cslAAV^20,000^ (cs refers cell-type-specific). Similarly, one of the two stranger rAAVs, rAAV2/9-EF1α- Flp, was diluted to ratios of 1:2,000 and 1:20,000 in PBS, followed by equal mixing with the other stranger rAAV, rAAV2/9-hSyn Con/Fon EYFP (the resulting rAAVs were abbreviated for cssAAV^2,000^ and cssAAV^20,000^, respectively).

For purpose (iii), the plasmids were AAV-EF1α-DIO-EGFP-TVA (GT) and AAV-EF1α-DIO-RG, and the littermate rAAVs, lAAV-DIO-GT/RG, had a ratio of 1:2. For the preparation of stranger rAAVs (sAAV-DIO-GT/RG), rAAV2/9-EF1α-DIO-GT and rAAV2/9-EF1α-DIO-RG were mixed at a ratio of 1:2.

For purpose (v), plasmids AAV-EF1α-EYFP and AAV-EF1α-mCherry were copackaged at a 1:1 ratio or independently packaged in HEK293T cells, generating the corresponding littermate rAAV2/9-EF1α-EYFP/mCherry (abbreviated for lAAV-EFYP/mCherry for convenience). As controls, the two stranger rAAVs, rAAV2/9- EF1α-EYFP and rAAV2/9-EF1α-mCherry, were mixed at a 1:1 ratio before experiments (abbreviated for sAAV-EFYP/mCherry).

The concentrations of the plasmids and the titers of all viruses are listed in **Supplementary Table 5.**

### Virus injections

All virus injection experiments were performed in Biosafety level 2 (BSL-2) environments. Mice were placed into a stereotaxic apparatus (#68030, RWD Life Science, China) and fixed with a nose clamp and ear bars after being deeply anesthetized by isoflurane. A small hole was made over the skull via a dental drill following the incision of the scalp along the midline of the brain. The coordinates for the injections were as follows based on *The Mouse Brain in Stereotaxic Coordinates*^65^: MOp (AP: 1.54 mm; ML: −1.6 mm; DV: −1.6 mm); CA3 (AP: −1.7 mm; ML: −2.0 mm; DV: −2.0 mm); CA1 (AP: −1.7 mm; ML: −1.0 mm; DV: −1.5 mm); basal forebrain: (AP: 0.21 mm; ML: −1.5 mm; DV: −5.5 mm) and VTA (AP: −3.4 mm; ML: −1.6 mm; DV: - 1.6 mm).

For comparisons of transsynaptic spreading efficiency (**Figs. 1,2** and **Supplementary Figs. 2-4**), the procedures for MAP-ENVIVIDERS and the classical rAAV-RV systems were similar to the reported^32, 33^, with injected volume as 80 nl for both lAAV-DIO-GT/RG and sAAV-DIO-GT/RG. Three weeks later, 150 nl EnvA-SADΔG-DsRed was injected into the same site of Thy1-Cre transgenic mice.

For non-cell-type-specific and cell-type-specific neuronal labeling, 100 nl littermate rAAVs (lAAVs and cslAAVs) and stranger rAAVs (sAAVs and cssAAVs) were injected for C57BL/6J mice (**Fig. 3,4** and **Supplementary Figs. 5-10**,**11c**,**14**) and Cre-driver transgenic lines (**Fig. 5,6** and **Supplementary Figs. 11a,b,d,e,12-15).**

For comparisons of sAAV-EFYP/mCherry with pure rAAVs (**Fig. 7a-c**) and sAAV-EFYP/mCherry with lAAV-EFYP/mCherry (**Fig. 7d-f**), similar titers of sAAV- EFYP/mCherry, rAAV2/9-EF1α-EYFP/PBS mixture (ratio of 1:1) and rAAV2/9- EF1α-mCherry/PBS mixture with a volume of 100 nl were injected into the MOp of C57BL/6J mice.

All virus solutions used were injected with a pulled glass micropipette at a rate of 20 nl/ min. The micropipette remained in place for at least 5 min before withdrawal from the brain. After recovery, the mice were housed carefully until perfusion.

### Perfusion and slicing

Three weeks after AAV injections and nine days after rabies virus injections, mice were perfused transcardially with 0.01 M PBS followed by 4% paraformaldehyde (PFA) in 0.01 M PBS. The extracted intact brains were postfixed overnight at 4°C and placed in 30% sucrose for 48-72 h, and 50- μm frozen sections were maintained across the whole brain (except that one mouse of lAAV^1,000,000^ where sections of injection site were cut into 100- μm to maintain the structures of local axons, **Supplementary Fig. 5a**) with a freezing microtome (CryoStar NX50 cryostat, Thermo Scientific, San Jose, CA). All brain slices were stored in a 24-well plate with PBS, and representative slices with labeled neurons in the injection site and long-range projections away from the injection site were identified with a Slide-scanning Microscope (Nikon Ni-E, Japan) for further analysis.

### Immunohistochemistry

Brain slices at the injection site of cslAAV^20,000^-labeled ChAT-Cre transgenic mice (**Fig. 5c** and **Supplementary Fig. 7**) and cslAAV^8,000^- labeled DAT-Cre transgenic mice were selected for immunostaining. The following antibodies: (i) Primary antibody, including goat anti-choline acetyltransferase (Millipore, AB144p, 1:200) and Rabbit anti-TH (abcam, ab112, 1:500); (ii) Secondary antibody, rabbit anti-Goat cy3 (Jackson Immunoresearch, 305-165-003, 1:500), Goat anti-rabbit Alexa 555 (Invitrogen, A21429, 1:500) were used in this study. Breifly, the selected 50-μm floating sections were initially blocked in 10% rabbit serum in PBS with 0.3% Triton X-100 (PBST) for 1 h at 37℃ and then washed with 0.3% PBST 10 min three times. Subsequently, sections were incubated in corresponding primary antibodies at 4℃ for 48-72 h followed by washes with 0.3% PBST 10 min three times and incubation in corresponding secondary antibodies at room temperature for 2 h.

### Image Acquisition

All brain slices were counterstained with 4′,6-diamidino-2- phenylindole (DAPI) to determine the cortical and laminar borders, coverslipped with Anti-fade Fluorescence Mounting Medium (Beyotime Biotechnology, China) and sealed with nail polish for imaging.

To quantify the whole-brain direct input neurons (**Fig. 2**, **Supplementary Fig. 4** and **Supplementary Table 1**), all coronal sections throughout ten brains labeled by MAP-ENVIVIDERS and classical rAAV-RV system were obtained through Nikon VS120 (Japan) with two (one for DAPI, one for DsRed) or three channels (the third for EGFP in injection site).

Except for images for input cells counting, all the other fluorescent images presented were acquired with a confocal laser scanning microscope (LSM710; Carl Zeiss, Germany) equipped with 405/488/514/561/633 excitation laser lines. 10x objectives (NA0.5) with a step size of 2 μm, 20x (NA0.8) with a step size of 1 μm, and 40x oil (NA1.4) with a step size of 0.39 μm were used. Generally, whole coronal sections containing the injection site were acquired at 10x (NA0.5) with a zoom of 1 or at 20x (NA0.8) with a zoom of 0.6. Regions of labeled somas were imaged and obtained in z-stacks with 10x (step size: 2 μm) or 20x objectives (step size: 1 μm). To obtain fine structures such as dendrite and axonal arborizations, 40x oil (step size: 0.39 μm) with a zoom of 1 was used for most images except for proximal spines of CA3 pyramidal neurons, where a zoom of 2 or 3 with the same objective was used (**Supplementary Fig. 8d**).

To faithfully compare the signal intensities among different labeling conditions (single virus vs. mixed viruses, littermate vs. stranger mixture, and littermate mixture vs. Thy1-EYFP-H line transgenic mice), parameters were adjusted low enough to ensure the fluorescent signal unsaturated, and then the same set of parameters was applied to all samples. The acquired z-stacks were stored in 16-bit depth TIFF format.

### Resin Embedding

For whole-brain imaging with the TDI-fMOST system, a lAAV^200,000^-labeled MOp mouse sample and a cslAAV^20,000^-labeled basal forebrain mouse sample were embedded with Technovit 9100 Methyl Methacrylate (MMA, Electron Microscopy Sciences, USA) according to previous procedures with minor optimizations^66^. Briefly, the removed PFA postfixed mouse brains were first rinsed in 0.01 M PBS for 12 h followed by complete dehydration in a series of alcohol (50%, 75%, 95%, 100%, and 100% ethanol, 2 h for each) and then xylene solution (twice, 2 h for each) for transparentization. The transparent brain was treated with sequential infiltration solutions (50%, 75%, 100%, and 100% resin in 100% ethanol, 2 h each for the first three solutions and 48 h for the final solution). Finally, the infiltrated brain was placed into a gelatin capsule filled with polymerization solution and kept in a dry chamber at −4°C in the dark for 72 h for polymerization before whole-brain sectioning.

### Dual-color, whole-brain imaging with the TDI-fMOST system

Brain-wide images of the embedded mouse brain were acquired with the TDI-fMOST system at a voxel size of 0.176 × 0.176 ×1 μm^3^ (unpublished data)^37^. Dual-color, whole-brain imaging with real-time propidium iodide (PI) staining was performed as previously described^67^. The embedded mouse brain was immersed in 0.05 M Na_2_CO_3_ solution to enhance the EYFP signal and decrease the fluorescence background during the process of imaging and sectioning^66^. A 60x water-immersed objective (NA 1.0) was used to image the surface layer of the brain sample with an axial step size of 1 μm each cycle prior to sectioning with a diamond knife. Each coronal section was acquired with 16-bit and 8-bit depth for the green and PI channels, respectively. After ten to twelve days of uninterrupted imaging and sectioning, we eventually obtained whole-brain raw data with a total of 12,841 continuous coronal sections for the MOp brain sample (70.2 TB with 46.4 TB for green and 23.8 TB for red channels) and 10,781 continuous coronal sections for the BF brain sample (54.7 TB with 36.8 TB for green and 17.9 TB red channels), respectively.

### Image processing

Raw data acquired with the dual-color TDI-fMOST system were preprocessed based on reported methods^67^, including seamless stitch and image registration for two channels. The preprocessed images were stored in an LZW compression TIFF format, with 16-bit depth for the green channel and 8-bit depth for the PI channel. For the generation of 3D reconstructions of brain-wide long-range projections of the MOp and basal forebrain mouse sample, we selected 12,740 and 10,780 continuous coronal sections from the respective green-channel data set and resampled to 2 × 2 × 4 μm^3^ before loading them into the Amira software (Visage Software, San Diego, CA). For the presentation of the MOp whole-brain dataset (**Supplementary Fig. 9**), coronal sections with maximum intensity projections (**Supplementary Fig. 9a-m**) were resampled to 0.6 × 0.6 μm^2^ due to the large image size, and images to display fine structures, including somas, axonal branches and axonal arborizations (residual images in **Supplementary Fig. 9**) were all selected from corresponding coronal sections of original resolution (0.176 × 0.176 μm^2^).

Raw data acquired with Nikon VS120 for whole-brain cell counting of input networks were transformed into JPG formats suitable for the following analysis. All confocal images were stored in lossless TIFF format. Image processing, including maximum intensity projection of serial z-stacks, adjusting brightness, and contrast of the composite images and selection of regions of interest, were performed with ImageJ (National Institutes of Health, USA). In addition, images imported from the Amira software were also processed as mentioned.

### Morphology reconstructions of individual neurons in MOp and BF

These reconstructions were performed in the Filament Editor module of the Amira software (Mercury Computer Systems, San Diego, CA, USA) via a human-machine interaction pattern. For the reconstructions of neurons 1-8, we randomly chose two neurons in L2, one neuron in L3, one neuron in L6a and four neurons in L5b and started neuronal tracing from the soma. We first traced local axons projecting from the main axons, followed by the tracing of main axons projecting to contralateral parts or subcortical regions. For the reconstructions of neurons **s1-s3**, we selected axons in the anterior commissure (ac) as a start point and traced in two opposite directions, with one towards the ipsilateral part and the other towards the contralateral part. For the reconstructions of neurons **c1-c4**, we selected complex axonal arborizations in the ipsilateral anterolateral/lateral visual area (VISal/l), ipsilateral caudoputamen (CP), ipsilateral and contralateral hippocampus as the start point and traced retrogradely. Because the axonal arborizations of most neurons were extremely complex and intermingled with each other, we made detailed registrations of the number and projection directions of all the branches to avoid negligence. For better differentiation, each part of the neuron, including the local axons, ipsilateral branches, and contralateral branches, were reconstructed separately. A constant 1000 × 1000 × 400 μm^3^ data block was imported into the Amira software in each cycle of tracing. After complete reconstructions of all arborizations of one branch, we moved to the next branch and repeated the tracing steps until all the branches projecting from the main axons were reconstructed. During the process of tracing, two tracing modules, thin structure and linear options were used alternatively according to the sparseness of the axonal projections. For example, if the axonal projections were sparse and not interfered by other axons, the thin structure module was used to depict the axonal projection path from the start point to the end; otherwise, the linear module was utilized to gradually trace the path. The brightness and thickness of the data block were adjusted from time to time in the process of tracing based on the different conditions of the axonal projections. The dendrite morphologies of all neurons were reconstructed in the same way as axonal tracing but separately for convenient subsequent analysis. The tracing results for each neuron, including the axonal projections and dendrite morphologies, were rechecked by different skilled technicians, and neurons with uncertainties were excluded from the final results. The locations of nuclei were based on the Allen Brain Atlas (http://mouse.brain-map.org/static/atlas) with the aid of PI-staining signals.

The reconstructed axons and dendrites of neurons **s1-s3** and **c1-c4** were saved in SWC files for the quantitative analysis of the branches and length in the software Neurolucida Explorer (MicroBrightField, USA).

### Axonal bouton counting

To make faithful manual counting of axonal boutons, we initially processed the raw signal of cslAAV^20,000^-labeled basal forebrain mouse sample with 20- to 100 μm-thickness maximum-intensity projections depend on the sparseness of axons. Subsequently, we randomly selected ten regions of interest (ROIs) from respective coronal sections of four cholinergic neurons (No. c1-c4). Counting of axonal boutons along the axonal shaft in each ROI were performed manually using multi-point tool in ImageJ according to the previously published cirteria^15, 16^ (**Figs. 6d** and **Supplementary Fig. 15**). The axonal pathway in each ROI was depicted in Amira software as mentioned above. The averaged bouton density (bouton/μm) in each ROI was defined as the ratio of total number of boutons over axonal length (**Figs. 6g** left). Based on the averaged bouton density of ten ROIs and total axonal length of reconstructed neurons, estimated total number of boutons of each cholinergic neuron was thus obtained (**Figs. 6g** right).

### Statistical analysis

Measurements of the signal profile (**Figs. 7a,b,e** and **Supplementary Figs. 4a**,**7b**) and relative fluorescence intensity (**1d**,**2f**,**3d**,**e**,**5i**,**6c**,**f** and **Supplementary Figs. 1**) were based on previously published methods with ImageJ^31^. Briefly, the signal profile of each pixel was calculated by subtracting the total signal value from the mean gray value of the background. We first measured the total signal value by a straight line drawn across the signal area (somas in the injection site or long-range axons) followed by measuring the mean gray value of the background by another straight line drawn across the background area close to the signal spot. For the measurement of relative fluorescence intensity, a rectangular selection (white dashed boxes) covering the signal spot (neuronal cell bodies) was made followed by the recording of the area of the boxes and the integrated density (IntDen) of the signal. To calculate the mean background, a straight line (yellow lines) was drawn across the background area close to the signal spot, and the mean gray value of the background was measured. The relative fluorescence intensity of each pixel was calculated by the relative signal values (resulting from subtraction of the total signal value from the total gray value of the background) over the pixel number of the area.

For comparison of the ratio between the signal intensity of DsRed and EGFP in starter cells labeled by lAAV-DIO-GT/RG and sAAV-DIO-GT/RG, a total of 170 starter neurons were selected randomly from 3 mice for both groups (**Fig. 1c** and **Supplementary Fig. 1a**, left panel. For comparisons of the signal intensity of DsRed in input cells within the lAAV-DIO-GT/RG- and sAAV-DIO-GT/RG labeling groups (**Fig. 2f** and **Supplementary Fig. 1a**, right panel), 160 input neurons for RT, 190 input neurons for SSp and SSs were selected randomly from 3 independent experimental mice for both groups.

For comparisons of the signal intensity of sparsely labeled neurons with non-cell-type-specific in wild-type mice (**Fig. 2d,e** and **Supplementary Fig. 1b,c**), 40 neurons from 2-5 brain slice of the injection site were selected randomly from three independent experimental mice for lAAV^20,000^, lAAV^200,000^, sAAV^20,000^, sAAV^40,000^ and sAAV^80,000^, and 20 out of 24 neurons were selected from lAAV^1,000,000,^ and 17 out of 26 neurons were selected from sAAV^1,000,000^ (the other 9 neurons were hardly captured under low imaging parameters). As comparisons, 40 neurons of MOp, CA1 and CA3 regions were selected randomly from brain slices of Thy1-EYFP-H transgenic mice with similar bregma.

For comparisons of the signal intensity of sparsely labeled cell-type-specific neurons (**Fig. 5i**), 40 neurons from 2-4 brain slices of the injection site were selected randomly from three independent cssAAV^20,000^-labeled Thy1-Cre mice, cssAAV^20,000^- labeled ChAT-Cre mice and cssAAV^8,000^-labeled DAT-Cre mice.

For comparisons of signal intensity of EYFP and mCherry (**Fig. 7c**), 90 neurons from brain slices of the injection site were selected randomly from 3 independent experimental mice for the sAAV-EYFP/mCherry, rAAV2/9-EF1α-EYFP/PBS mixture (ratio of 1:1) and rAAV2/9-EF1α-mCherry/PBS mixture (ratio of 1:1).

Whole-brain counting of input cells was performed following previously published methods^32, 33^. Briefly, input cells in each coronal section were identified manually, followed by registration of each coronal section to the Allen Brain Atlas (ABA, http://mouse.brain-map.org/). We summed the total number of input neurons and calculated the convergent index (ratio between input cells and starter cells)^8^ in diverse gross regions and subregions. For the demonstration of relationship between proportion of inputs and enhanced ratios of tracing efficiency provided by MAP-ENVIVIDERS (**Fig. 2e**), we calculated selected 44 representative subregions from 11 major regions from sAAV-DIO-GT/RG labeling groups and calculated proportion of inputs, i.e. inputs in these subregions against total inputs (for better demonstration, proportion of inputs in x-axis were listed by logarithm to base 10, denoted by lg %). Enhanced ratios of tracing efficiency provided by MAP-ENVIVIDERS (y-axis) were denoted by convergent index of lAAV-DIO-GT/RG dividing by convergent index of sAAV-DIO-GT/RG in diverse subregions.

All quantitative data, including cell numbers and measurement of relative signal intensity, convergent index and proportion of inputs, were all presented as the mean ± s.e.m. Significance was analyzed using one-way ANOVA with Dunnett’s post hoc test (**Fig. 3d**), Turkey’s post hoc test (**Fig. 5i**), Student’s t-test (**Figs. 1d,2f,3e,7c,f**) and Mann-Whitney U test (**Figs. 2a-d** and **Supplementary Fig. 4**) in GraphPad Prism version 6.0 (GraphPad Software Inc., San Diego, CA, USA).

## Supplementary Information

**Supplementary figure 1.**
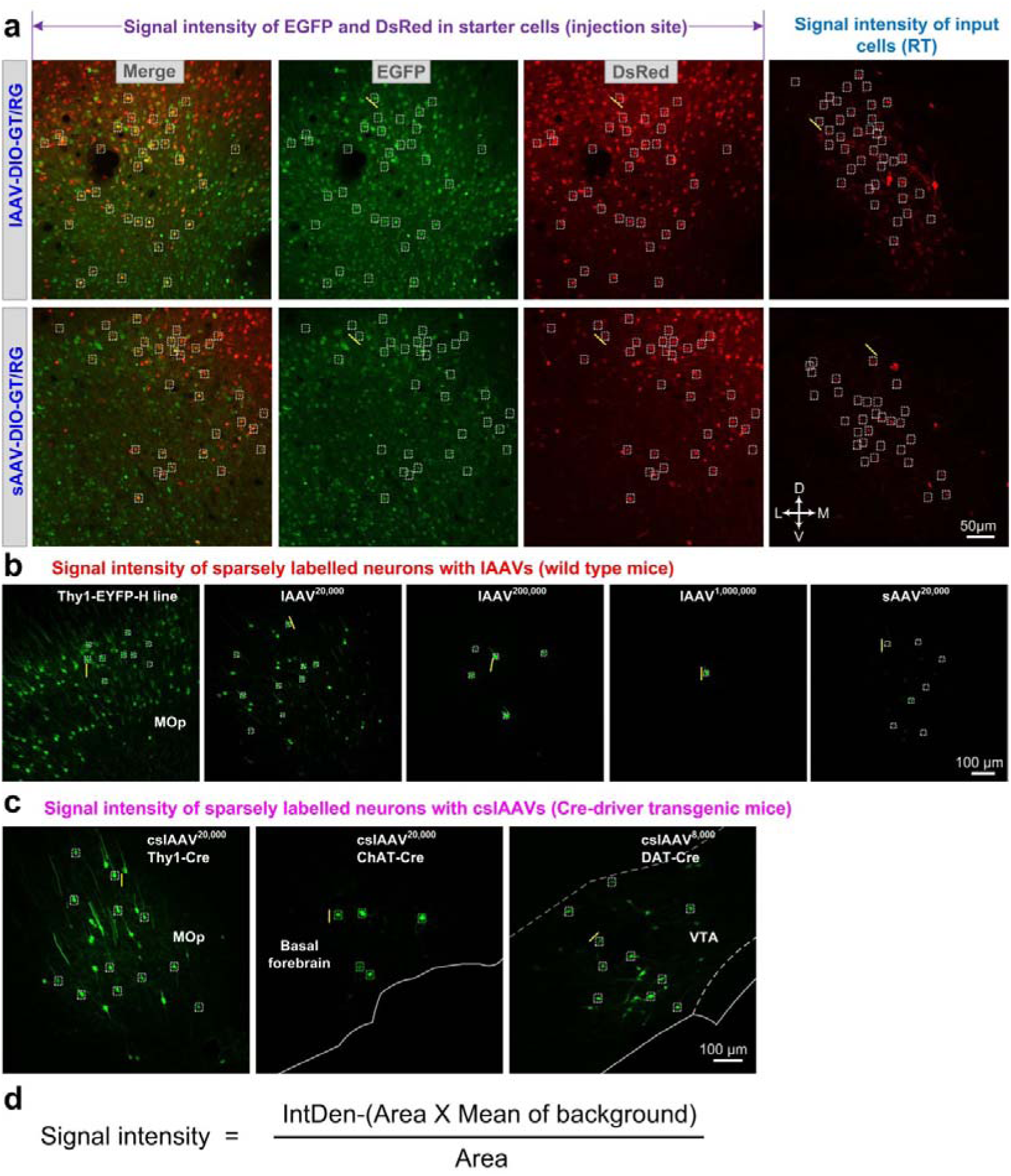
Schematic of measuring fluorescence intensity of individual neurons. **(a)** Representative confocal images showing the measurement of signal intensity of EGFP and DsRed in starter cells and DsRed-labeled input cells in lAAV-DIO- GT/RG- and sAAV-DIO-GT/RG-treated mice. All of the images were obtained under the same imaging conditions (low laser intensity was used to avoid over-exposure of the cell bodies). **(b)** Representative confocal images showing the measurement of signal intensity of neurons in Thy1-EYFP-H line transgenic mice and sparsely labeled neurons with diverse lAAVs and sAAVs in MOp region. **(c)** Representative confocal images showing the measurement of signal intensity of neurons in cssAAV^20,000^-labeled Thy1-Cre mice (MOp), cssAAV^20,000^-labeled ChAT- Cre mice (basal forebrain) and cssAAV^8,000^-labeled DAT-Cre mice (VTA). **(d)** The signal intensities were calculated in ImageJ according to the formula in **d** and the method in the previous report^31^. Briefly, a rectangular selection (white dashed boxes) covering the signal spot (neuronal cell bodies) was made, and then the area of the boxes and the integrated density (IntDen) of the signal were recorded. To calculate the mean background, a straight line (yellow lines) was drawn across the background area close to the signal spot and the mean gray value of the background was measured. This method was applied to calculate the signal intensities in **Figs 1d**,**2d**,**3d**,**e**,**5j**,**7c**. Abbreviations: MOp, primary motor area; RT, reticular nucleus of the thalamus; VTA, ventral tegmental area.

**Supplementary figure 2.**
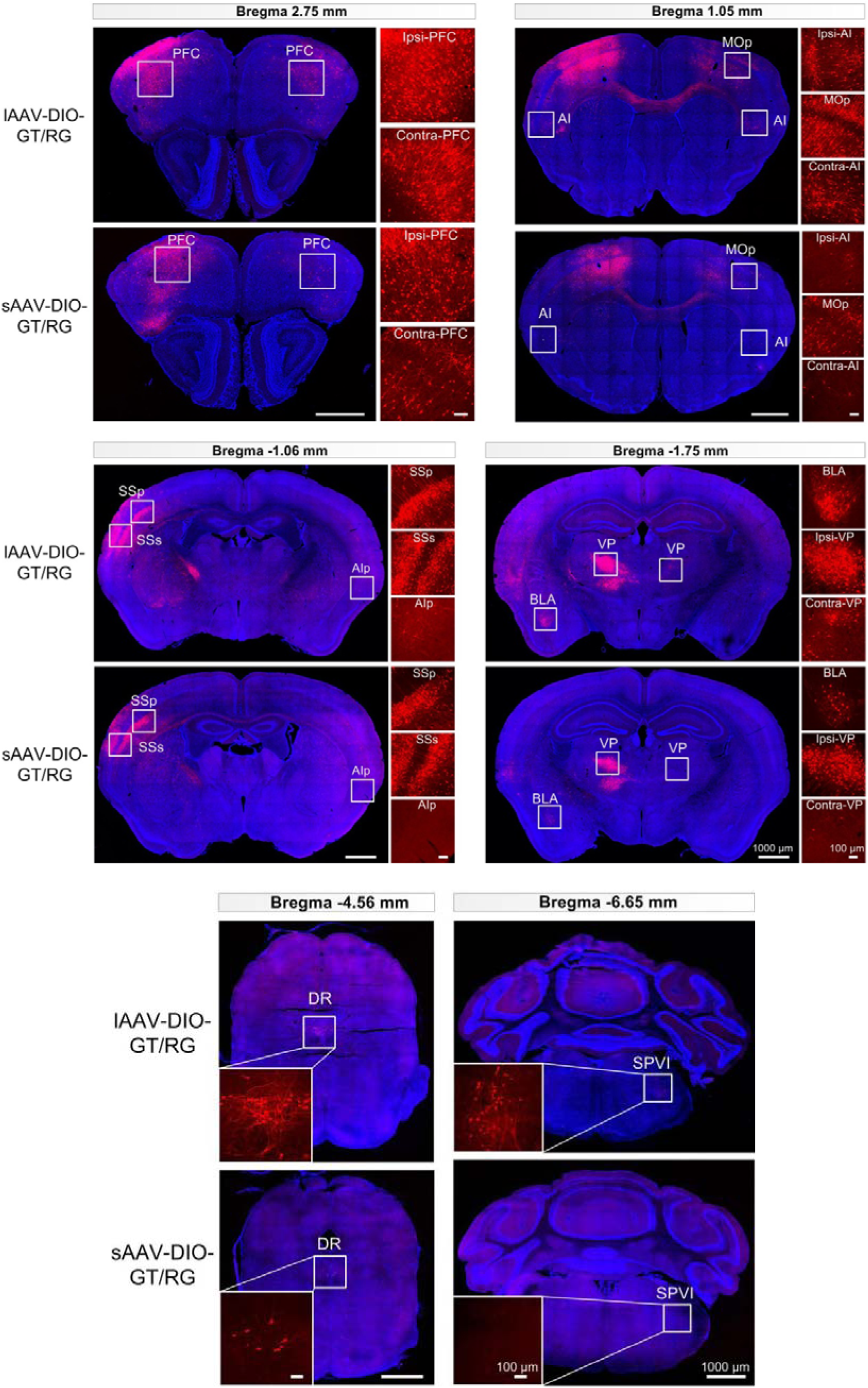
Localization of nuclei demonstrated in Fig. 6e with the aid of DAPI (4’,6- diamidino-2-phenylindole) staining. Abbreviations: AI, agranular insular area; AIp; agranular insular area, posterior part; BLA, basolateral amygdalar nucleus; DR, dorsal nucleus raphe; MOp, primary motor area; PFC, prefrontal cortex; SPVI, spinal nucleus of the trigeminal, interpolar part; SSp, primary somatosensory area; SSs, supplemental somatosensory area; VAL, ventral anterior-lateral complex of the thalamus; VP, ventral posterior complex of the thalamus.

**Supplementary figure 3.**
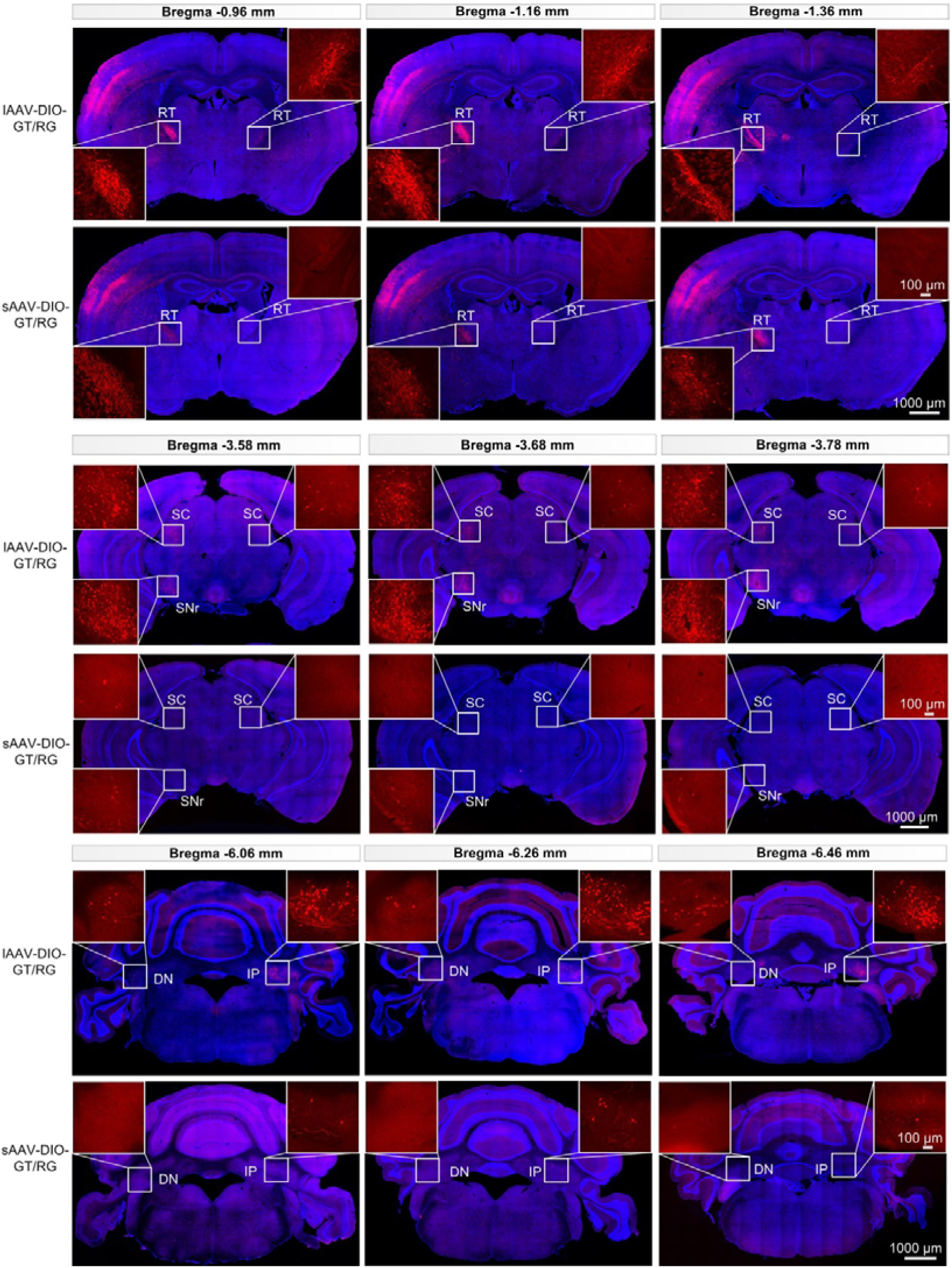
Comparisons of inputs in RT of thalamus, SC and SNr of midbrain, DN and IP of cerebellum between lAAV-DIO-GT/RG and sAAV-DIO-GT/RG labeling groups. Insets were enlargements respective brain regions. DN, dentate nucleus; IP, interpeduncular nucleus; RT, reticular nucleus of the thalamus; SC, superior colliculus; SNr, substantia nigra pars reticulate.

**Supplementary figure 4.**
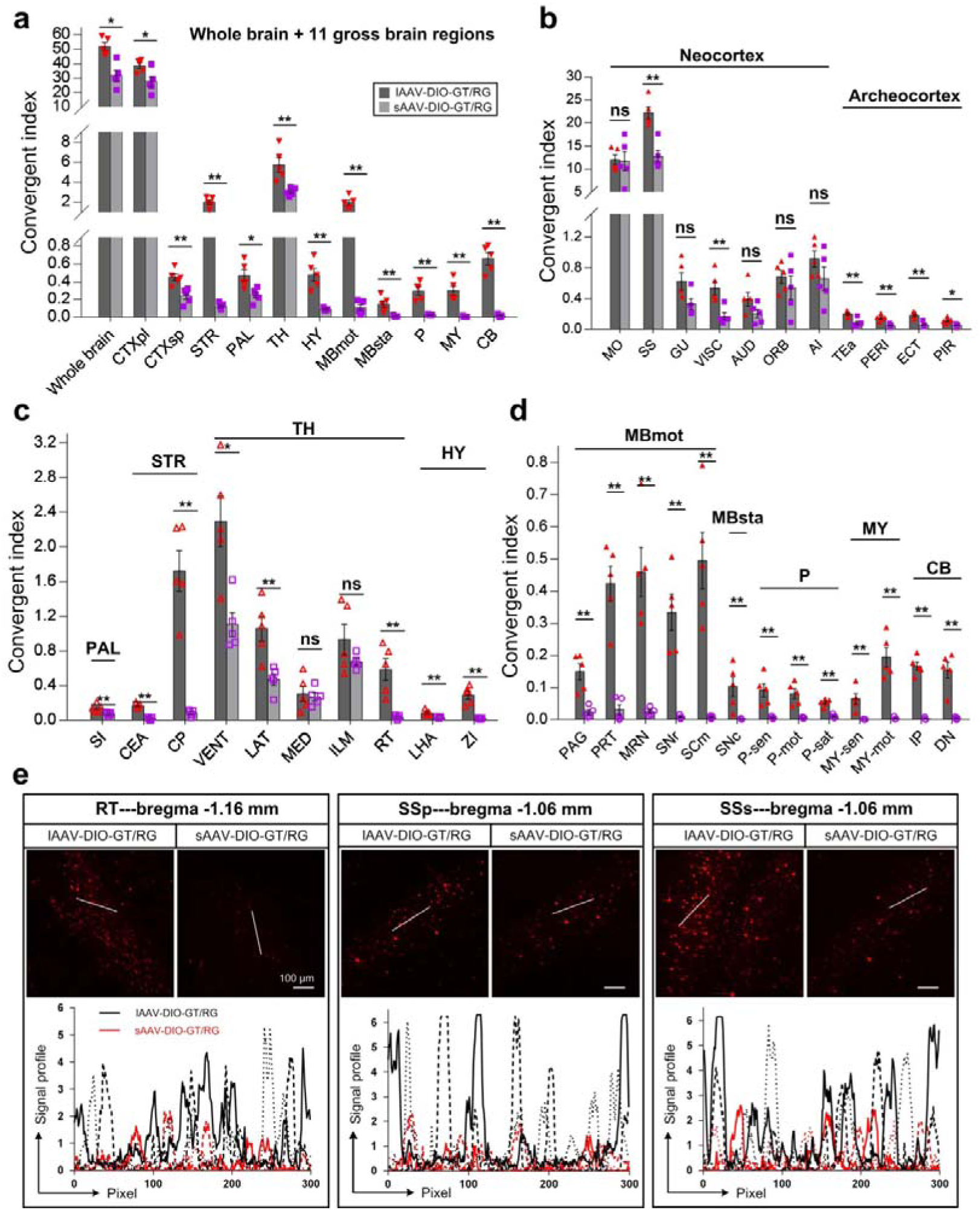
(a-d) Brain-wide comparisons of tracing efficiency between lAAV-DIO-GT/RG and sAAV-DIO-GT/RG labeling groups. **(a)** Comparisons of convergent index of total inputs and 11 gross brain regions. **(b)** Comparisons of convergent index in diverse cortical regions. **(c)** Comparisons of convergent index in pallidum (PAL), striatum (STR), thalamus (TH) and hypothalamus (HY). **(d)** Comparisons of convergent index in midbrain, motor related (MBmot), midbrain, behavioral state related (MBsta), pons (P), medulla (MY) and cerebellum (CB). All data are presented as mean ± s.e.m. n = 5 mice for each group in **a**-**d**. Mann-Whitney U test. n.s, non-significant (P > 0.05), *P < 0.05, **P < 0.01. **(e)** Signal profile of input neurons in RT, SSp and SSs in lAAV-DIO-GT/RG and sAAV-DIO-GT/RG labeling groups. n = 3 mice for each group. Abbreviations see **Supplementary Table 2**.

**Supplementary figure 5.**
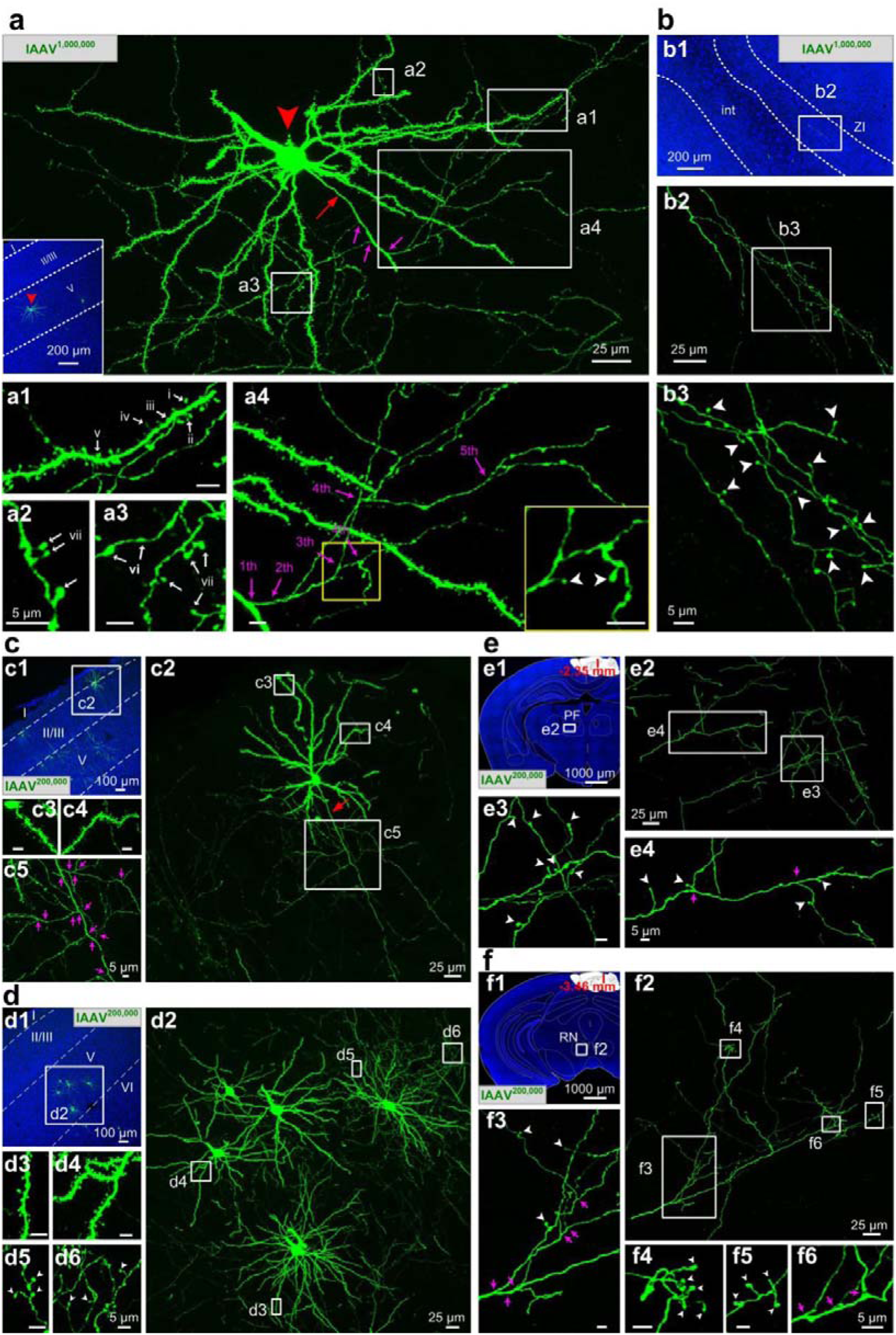
Fine structures of individual neurons labeled with lAAV^1,000,000^ and lAAV^200,000^. **(a)** Confocal image (100-µm thickness) of a representative individual neuron labeled with lAAV^1,000,000^ in injection site. Inset showed the localization of cell body (indicated by red arrowhead). **(a1)** Five types of spines: (i) mushroom, (ii) thin, (iii) stubby, (iv) filopodium, and (v) branched spines. **(a2-a3)** Two types of boutons: (vi) en passant boutons and (vii) terminal boutons. **(a4)** Different levels of local axonal collaterals. **(b)** Confocal images of representative long-range axonal projections in the ZI region (**b1**) labeled with lAAV^1,000,000^. Fine structures were progressively enlarged from **b1** to **b3**. **(c-d)** Fine structures of MOp L2 (**c**) and L5 (**d**) pyramidal neurons labeled with lAAV^200,000^ in injection site. **c1** and **d1** were low-magnification images indicating the locations of cell bodies. **c2** and **d2** were high-magnification images showing the fine structures of the representative neurons. **c3-c5** and **d3-d6** were enlargements of box regions in **c2** and **d2**, respectively. **(e-f)** Fine structures of MOp long-range axonal projections labeled with lAAV^200,000^. **e1** and **f1** were composite coronal sections indicated the locations of the axonal projections, including parafascicular nucleus (PF, **e1**) in thalamus and red nucleus (RN, **f1**) in midbrain. **e2** and **f2** were high-magnification images showing the fine structures of the complex axonal arborizations. **e3-e4** and **f3-f6** were enlargements of the boxed regions in **e2** and **f2**, respectively. Red arrows indicated the primary axon emitting from soma. The pink arrows and numerals indicate the orders of the axonal branches and bright, large bulb-shaped terminal boutons were indicated by white arrowheads. Definition the boundary of cortical layers (indicated by white dashed lines) and location of nuclei were all based on DAPI (4’,6-diamidino-2-phenylindole, blue) staining. int, internal capsule; ZI, zona incerta. I-VI, cortical layers I to VI.

**Supplementary figure 6.**
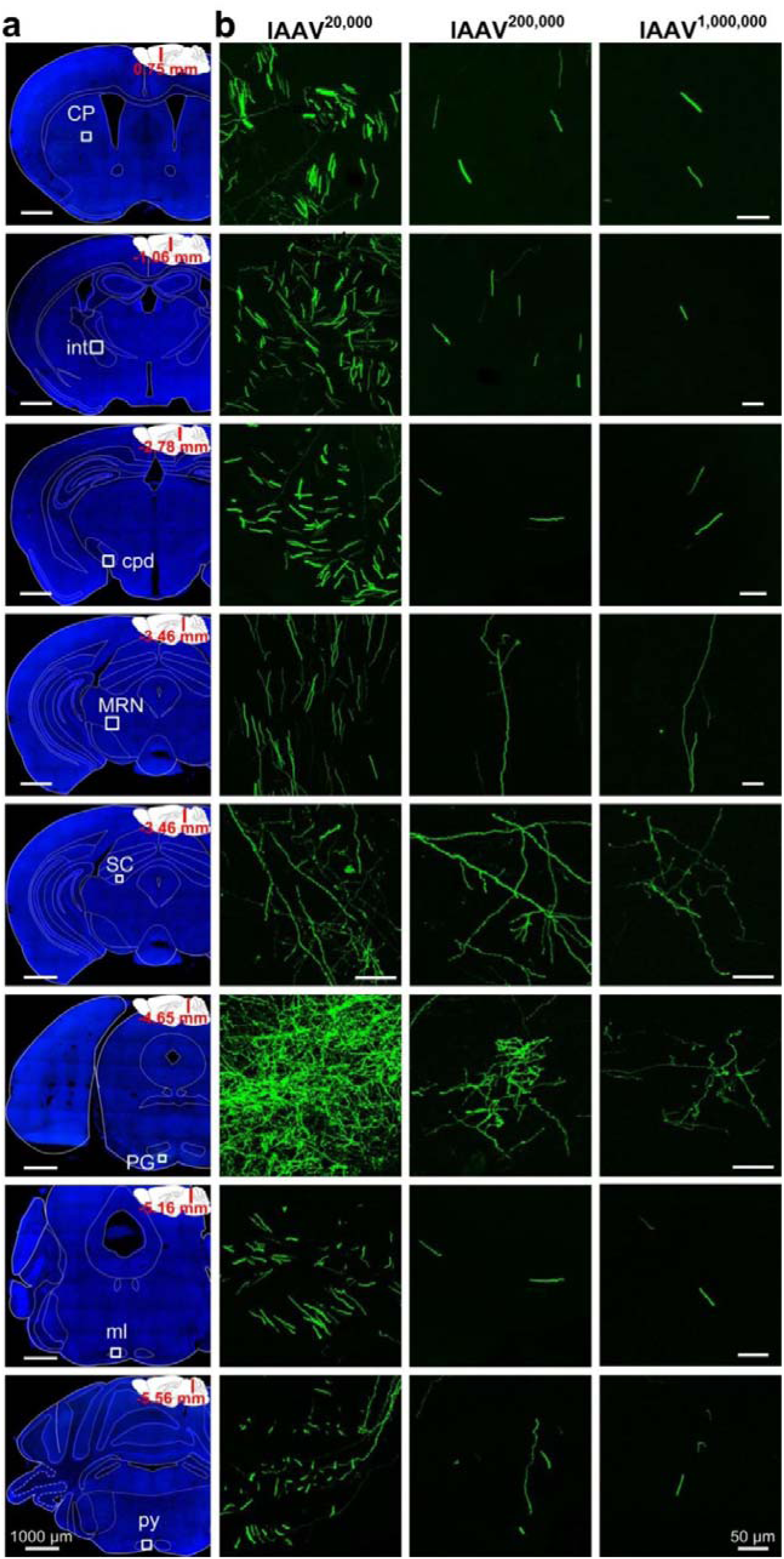
Brain-wide long-range axonal projections of three lAAVs in MOp. We injected equal volume (100 nl) of three lAAVs (lAAV^20,000^, lAAV^200,000^ and lAAV^1,000,000^) into MOp of wild-type mice. After 21d’s expression, we examined brain-wide long-range axonal projections of three lAAVs, including CP (bregma level: 0.75 mm), int (bregma level: −1.06 mm), cpd (bregma level: −2.78 mm), MRN and SC (bregma level: −3.46 mm), PG (bregma level: −4.65 mm), ml (bregma level: −5.16 mm), and py (bregma level: −5.56 mm). **(a)** Composite coronal sections showing the locations of the axonal projections. **(b)** Maximum-intensity projections of the boxed regions in **a**, showing the fine structures of the axonal projections in different brain regions. Images coming from the same brain region were obtained with the same imaging conditions and processed identically. Abbreviations: CP, caudoputamen; int, internal capsule; cpd, cerebal peduncle; ml, medial lemniscus; MRN, midbrain reticular nucleus; PG, pontine gray; py, pyramid; SC, superior colliculus.

**Supplementary figure 7.**
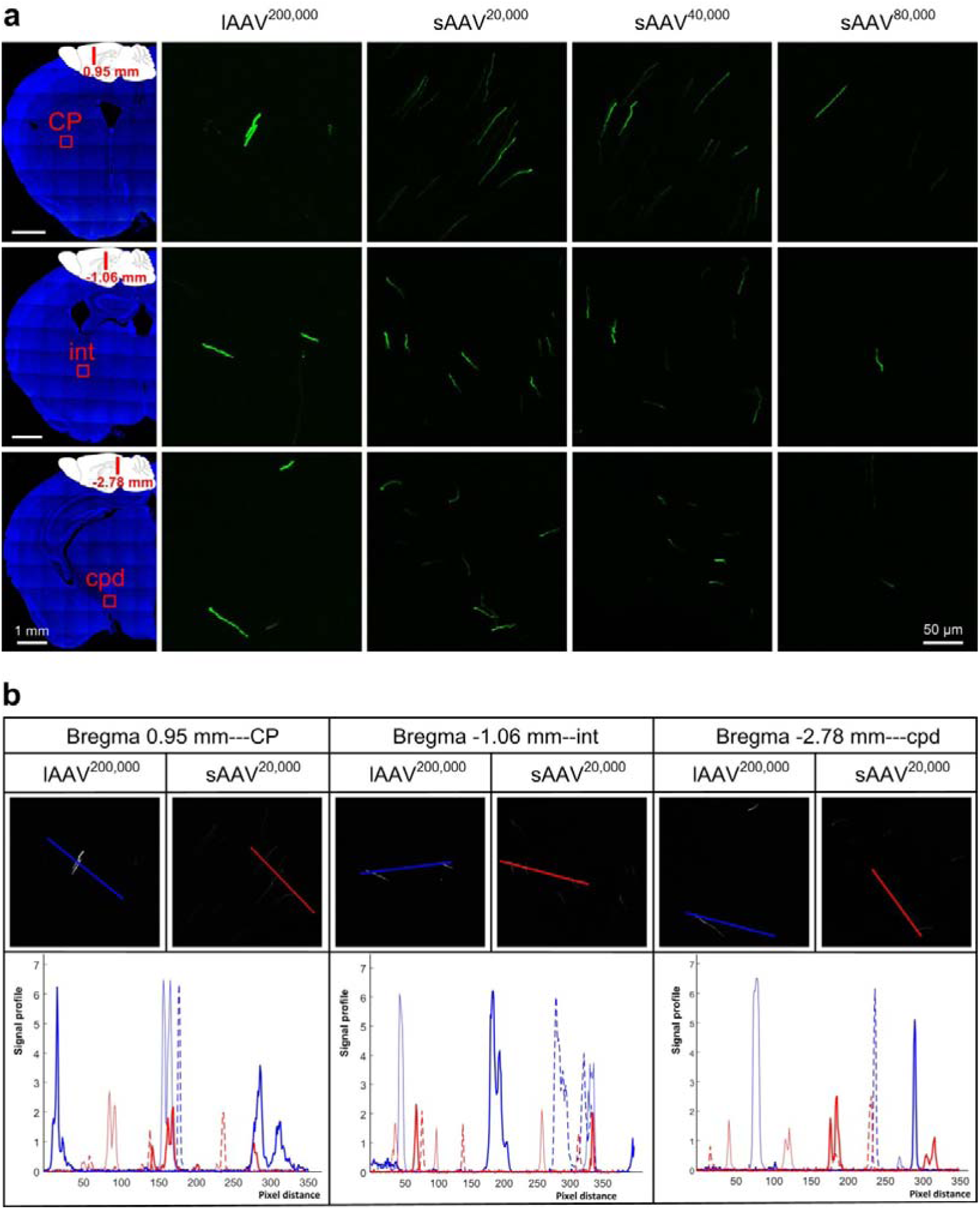
Comparisons of brightness of long-range axonal projections between lAAV^200,000^ and diverse sAAVs (sAAV^20,000^, sAAV^40,000^ and sAAV^80,000^). **(a)** Representative confocal images of long-range axonal projections in different brain regions. All images in each brain region were obtained under the same imaging conditions and processed identically. **(b)** Measurements of fluorescence intensities in **a** for lAAV^200,000^ (indicated by blue lines, three types of blue lines represent three different mice) and sAAV^20,000^ (indicated by red lines, three types of red lines represent three different mice). The data showed that the fluorescence in neurons labeled by lAAVs was stronger than by sAAVs.

**Supplementary figure 8.**
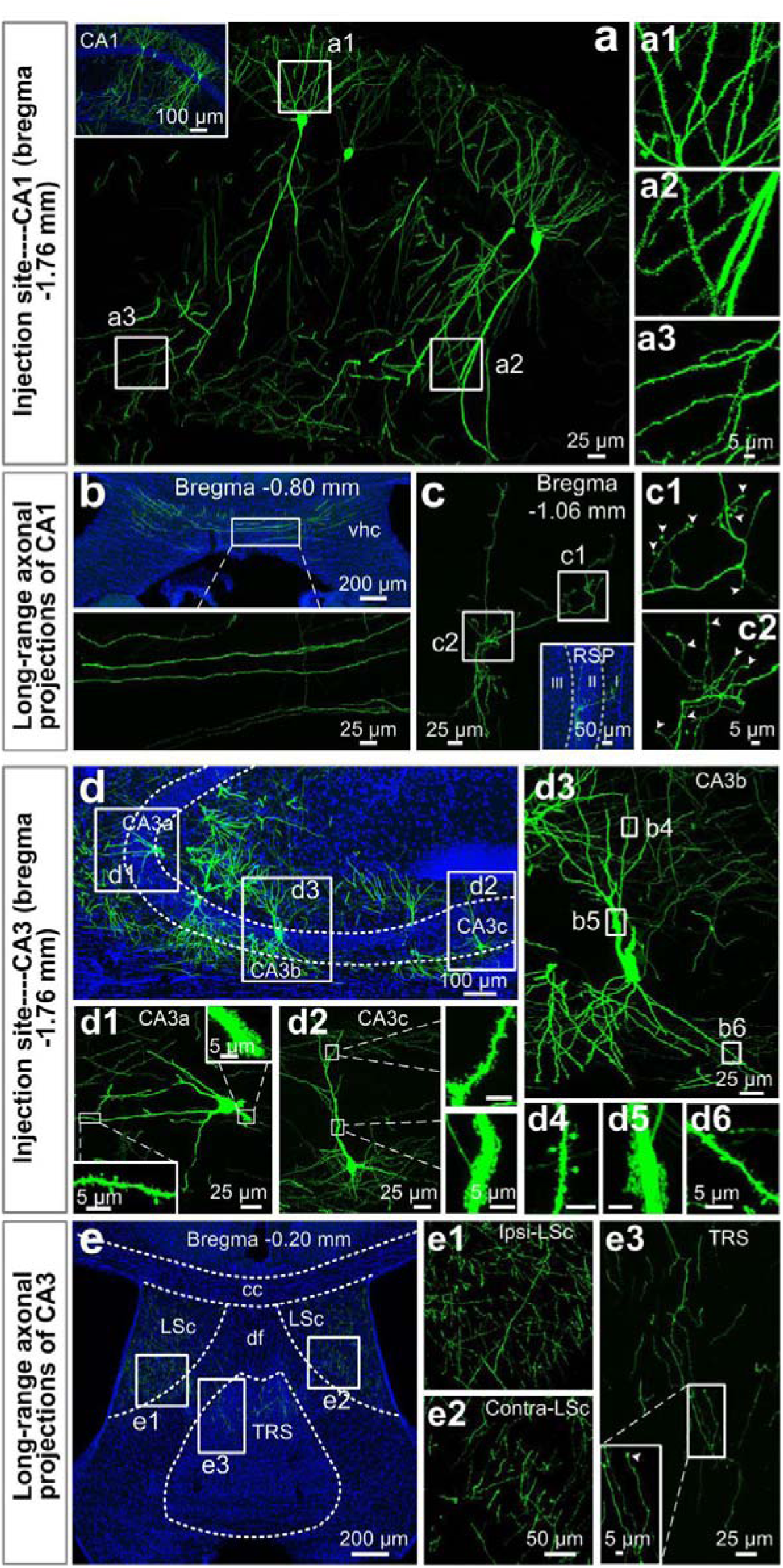
Applications of lAAV^200,000^ in CA1 and CA3 region. **(a-c)** Applications of lAAV^200,000^ in CA1 region. **(a)** Representative confocal images showing the fine structures of labeled neurons in injection site of CA1. Inset shows the locations of the somata. **(a1-a3)** Enlargements of the boxed regions in **a** showing different parts of dendritic spines. **(b-c)** Representative confocal images of the long-range axonal projections of CA1 in ventral hippocampal commissure (vhc, **b**) and retrosplenial cortex (RSP, **c**). **c1-c2** were enlargements of the corresponding boxed regions in **c**. Insets in **c** were approximate location of the axonal arborizations. White arrowheads indicate terminal boutons. White dashed lines on inset define the boundary of cortical layers based on DAPI staining. **(d-e)** Applications of lAAV^200,000^ in CA3 region. **(d)** Low-magnification images showing the overall labeling in CA3. **(d1-d3)** Fine structures of the CA3a (**d1**), CA3b (**d3**), and CA3c (**d2**) pyramidal neurons. Respective box regions in **d1** and **d2** were enlarged to show proximal and distal dendritic spines. **(d4-d6)** Enlargements of the box regions in **(d3)** showing apical **(d4),** proximal **(d5),** and basal dendritic spines **(d6)** of CA3b pyramidal neuron, respectively. **(e)** Low-magnification images indicate the locations of the axonal projections in TRS and LSc. The location of nuclei was based on Paxinos Mouse Brain Atlas (PMBA) with the aid of DAPI staining. **(e1-e3)** High-magnification images of the boxed regions in **e** showing the fine structures of axonal projections and axonal terminals. Abbreviations: CA1, field CA1 of hippocampus; CA3, field CA3 of hippocampus; cc: corpus callosum; df: dorsal fornix; DG, dentate gyrus; LSc: lateral septal nucleus, caudal (caudodorsal) part; TRS: triangular nucleus of septum.

**Supplementary figure 9.**
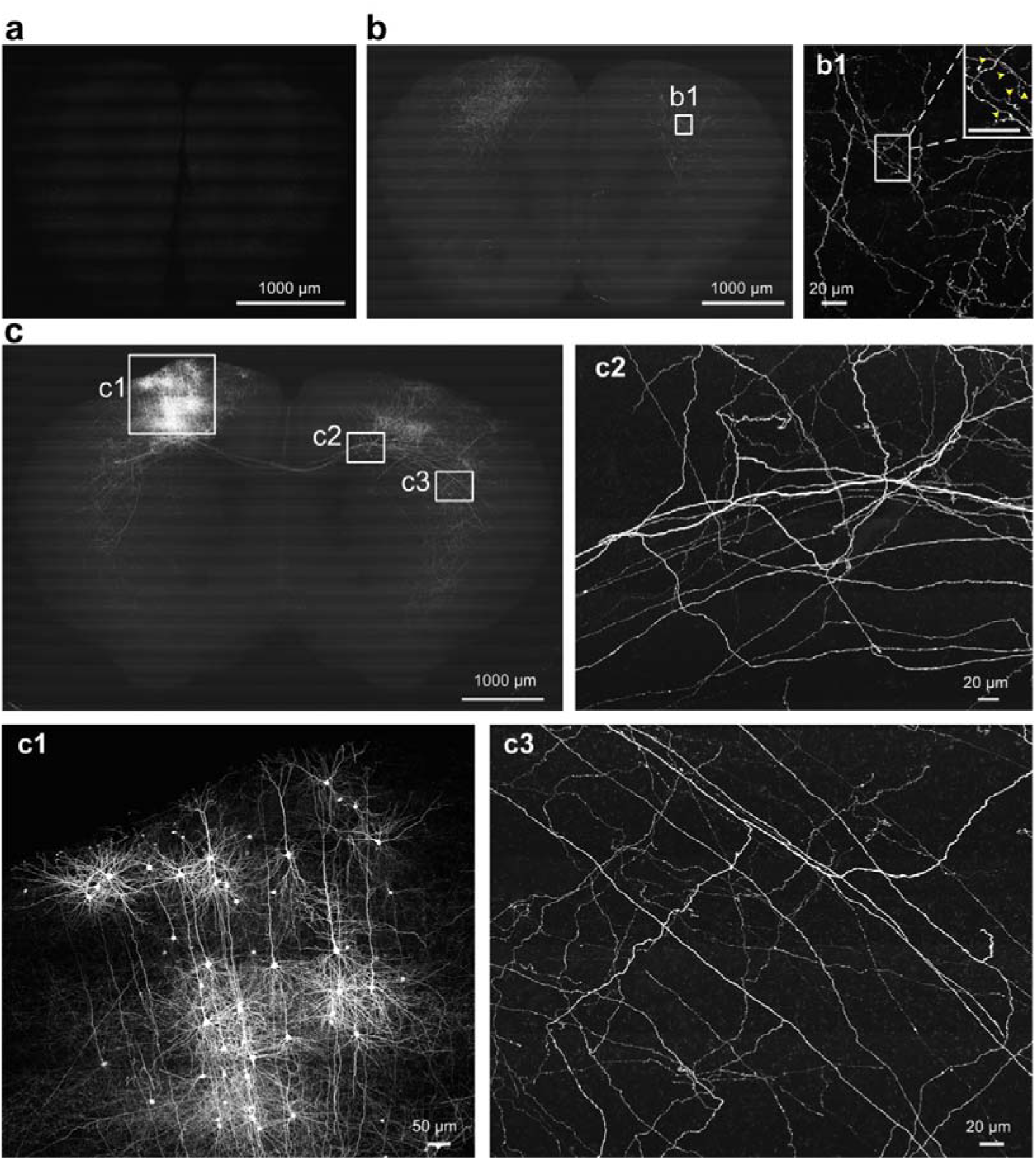
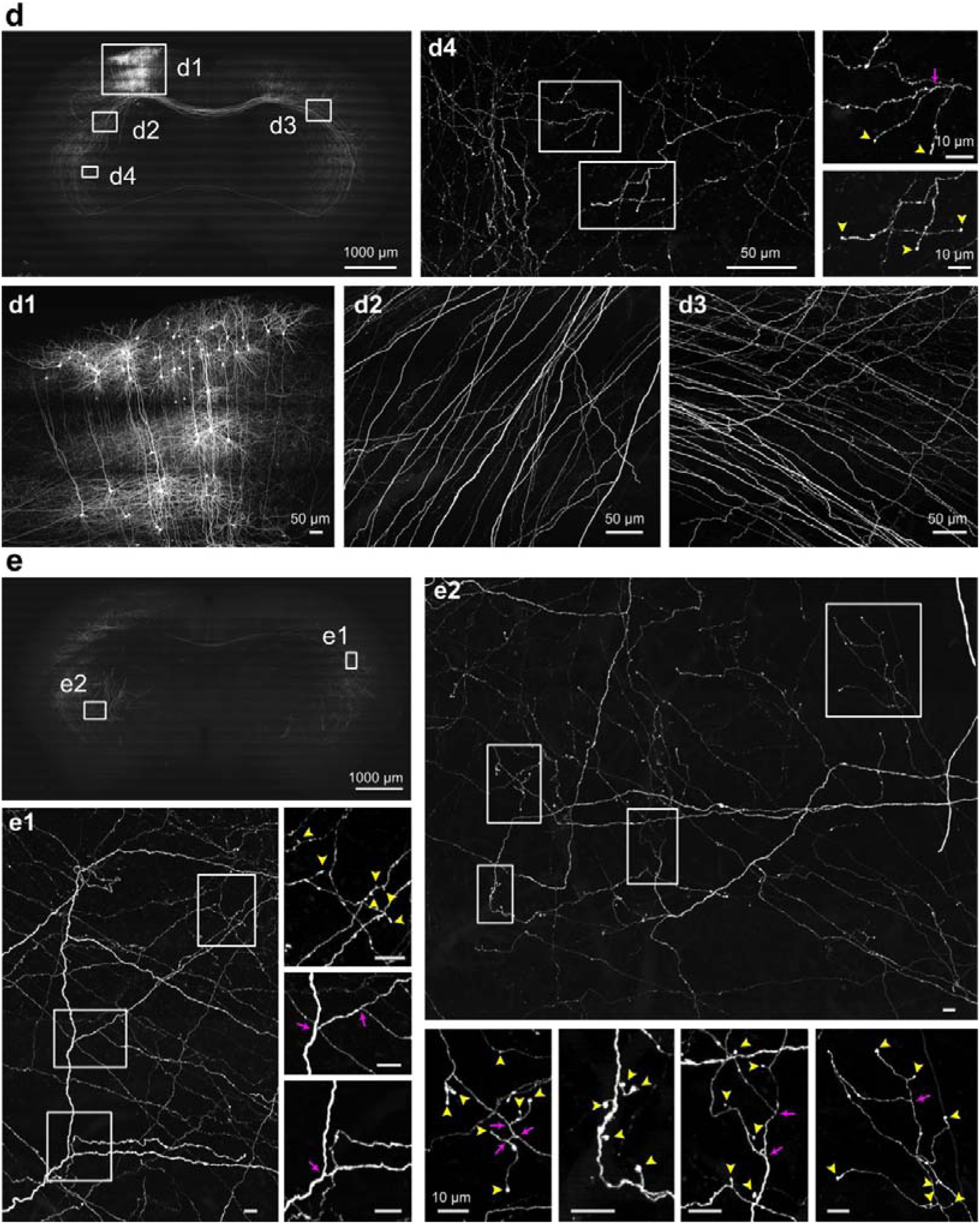
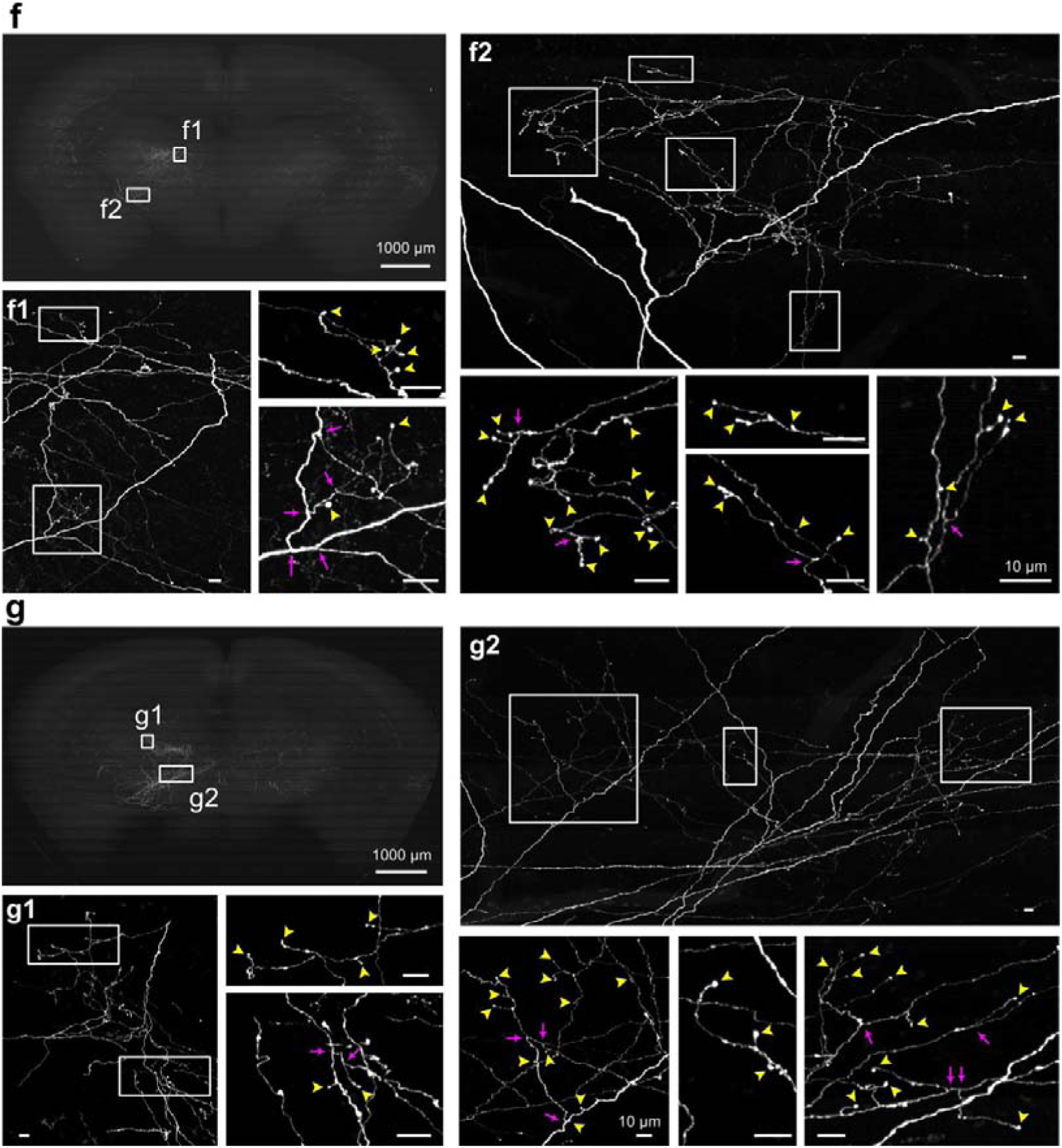
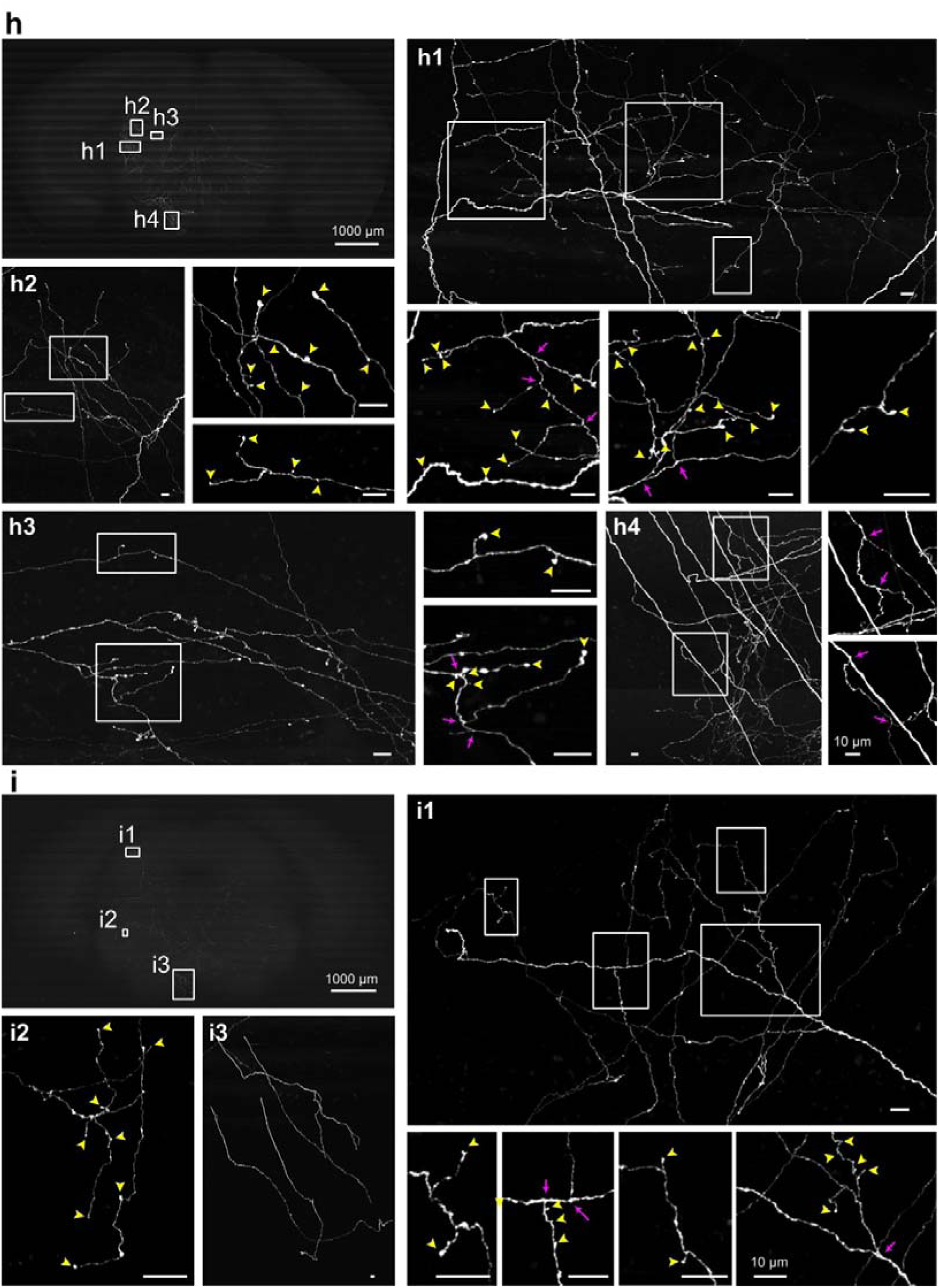
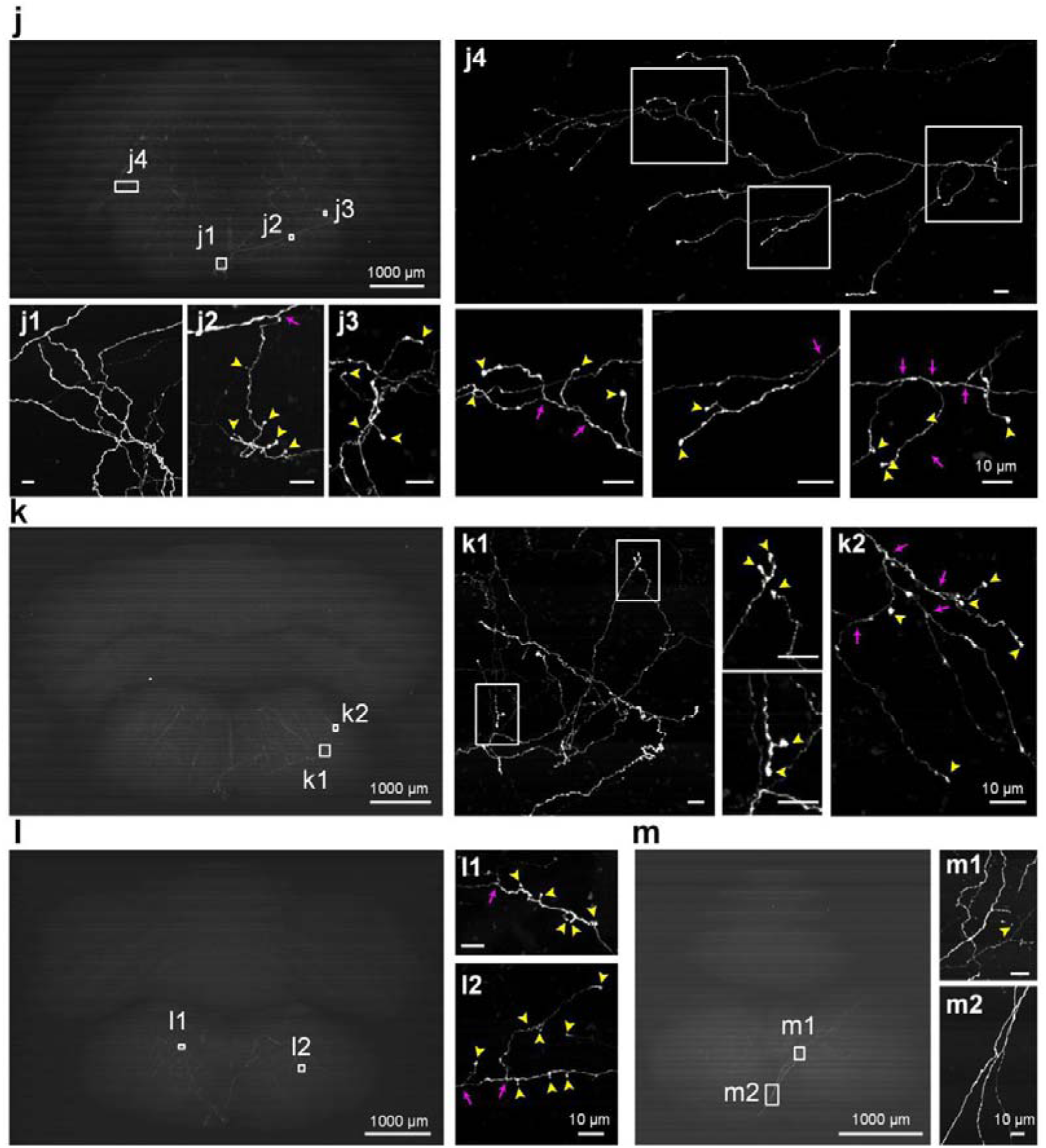
Whole-brain data sets of MOp (labeled with lAAV^200,000^). **a** and **b** were maximum-intensity projections of 800 and 1,600 coronal sections. **c-l** were maximum-intensity projections of 1000 coronal sections and **m** was maximum-intensity projections of 420 coronal sections, respectively. All other images were enlargements of respective box regions showing the injection site or fine structures, including axonal branches (indicated by purple arrows) and terminal boutons (indicated by yellow arrowheads).

**Supplementary figure 10.**
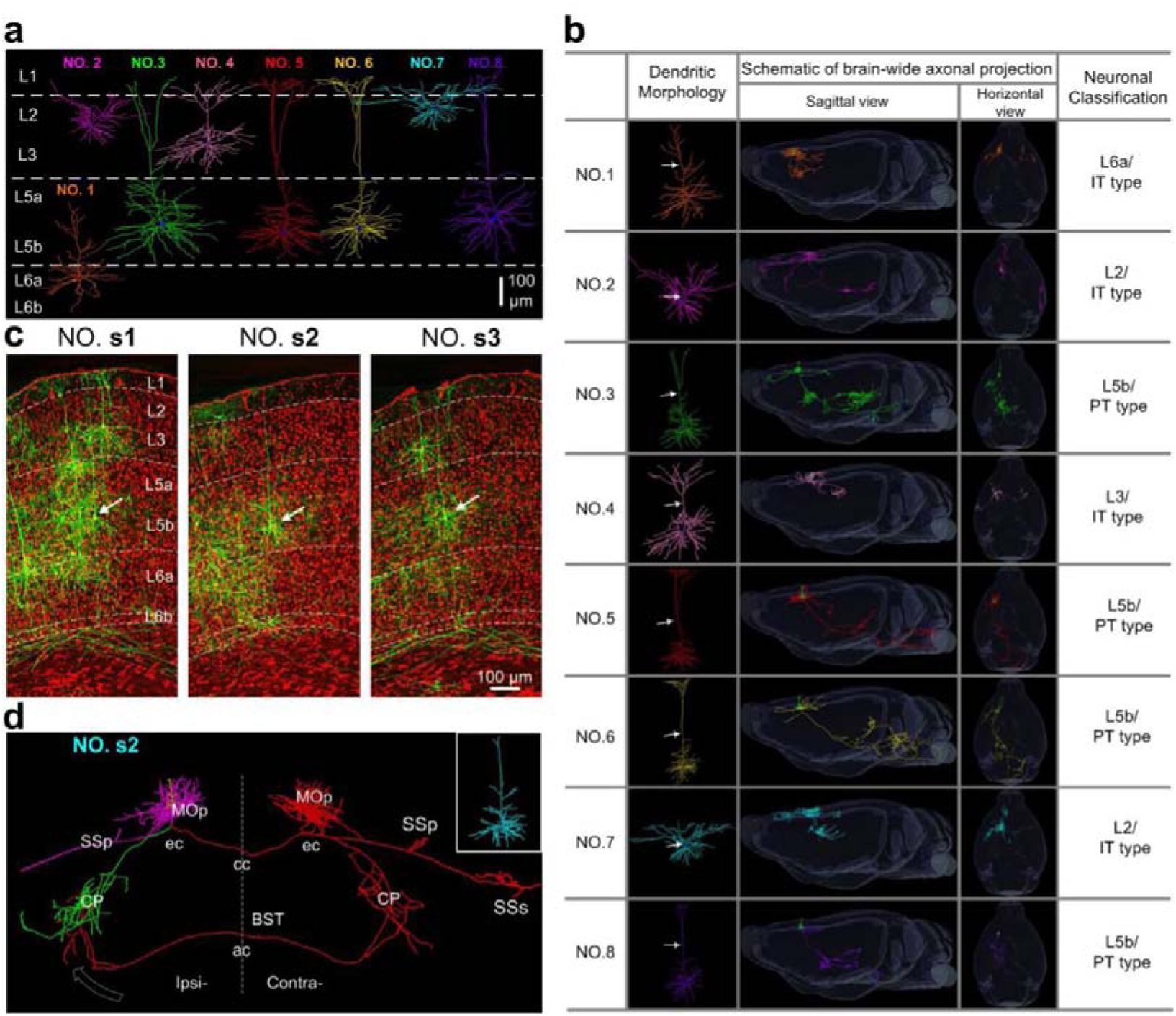
Localization, dendritic and axonal morphologies of the reconstructed MOp pyramidal neurons. **(a)** Gallery of dendritic morphologies and approximate laminar distributions of eight reconstructed pyramidal neurons (No.1-8). **(b)** Gallery of Axonal morphologies and probable classifications of 8 neurons. White arrows indicate apical dendrites. **(c)** Composite coronal sections show the locations of the cell body of neurons **s1**-**s3,** which were indicated by white arrows (EYFP signals were in green with 100-µm thickness and cytoarchitectonic reference, i.e. propidium iodide (PI) signals were in red with 10-µm thickness). White dashed lines define boundary of cortical layers. **(d)** Localization of the brain-wide axonal projections of neurons **s2** (dendrites: yellow; local axons: magnet; left branches and right branches originated from ipsilateral external capsule were in green and red color, respectively). The brain regions were defined according to the Allen Brain Atlas (ABA) with the aid of the PI signals. Dashed white arrows pointed terminated directions of main axons.

**Supplementary figure 11.**
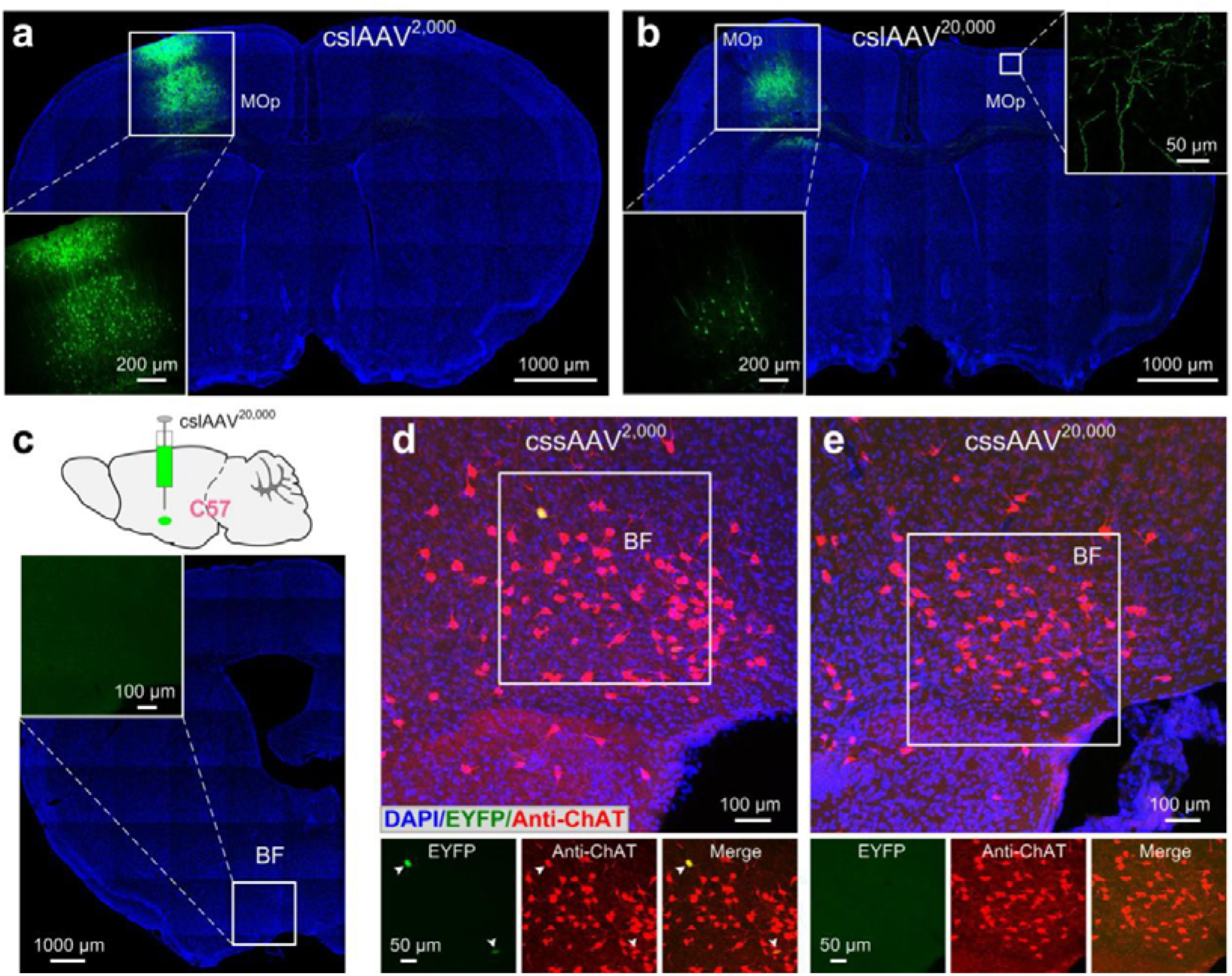
Validation of cell-type-specific labeling strategy with cslAAVs in Thy1-Cre and ChAT-Cre transgenic mice. **(a-b)** Two versions of cslAAVs, cslAAV^2,000^ (**a**) or cslAAV^20,000^ (**b**) were injected into MOp of Thy1-Cre transgenic mice to achieve tunable neuron labeling. Lower left insets in **a** and **b** were enlargements of the corresponding boxes showing labeling in injection sites. Upper right inset in **b** was contralateral MOp axonal projections. **(c)** 100nl cslAAV^20,000^ was injected into basal forebrain (BF) of wild-type mice, we found no neurons were labeled across the injection site of 4 injected mice 21d later, indicating that the expression of cslAAVs was strictly restricted to Cre-driver lines. **(d-e)** For control experiments, we injected 100 nl two diluted ratios of Flp-expressing rAAV: Flp- and Cre-dependent rAAVs mixtures (abbreviated for cssAAV^2,000^ and cssAAV^20,000^ hereafter, in which the Flp-expressing rAAV was diluted to ratios of 1:2,000 and 1:20,000 in PBS respectively before equally mixed with Flp- and Cre-dependent rAAV into BF of ChAT-Cre transgenic mice (n = 5 mice each). We found no labeled neurons were detectable in cssAAV^20,000^-treated mice and in cssAAV^2,000^- treated mice, only two mice each contained three or four faintly labeled cholinergic neurons (note that the imaging parameters of EYFP were much higher than Fig. 3c) whereas the other three mice rarely contained any signals in injection sites.

**Supplementary figure 12.**
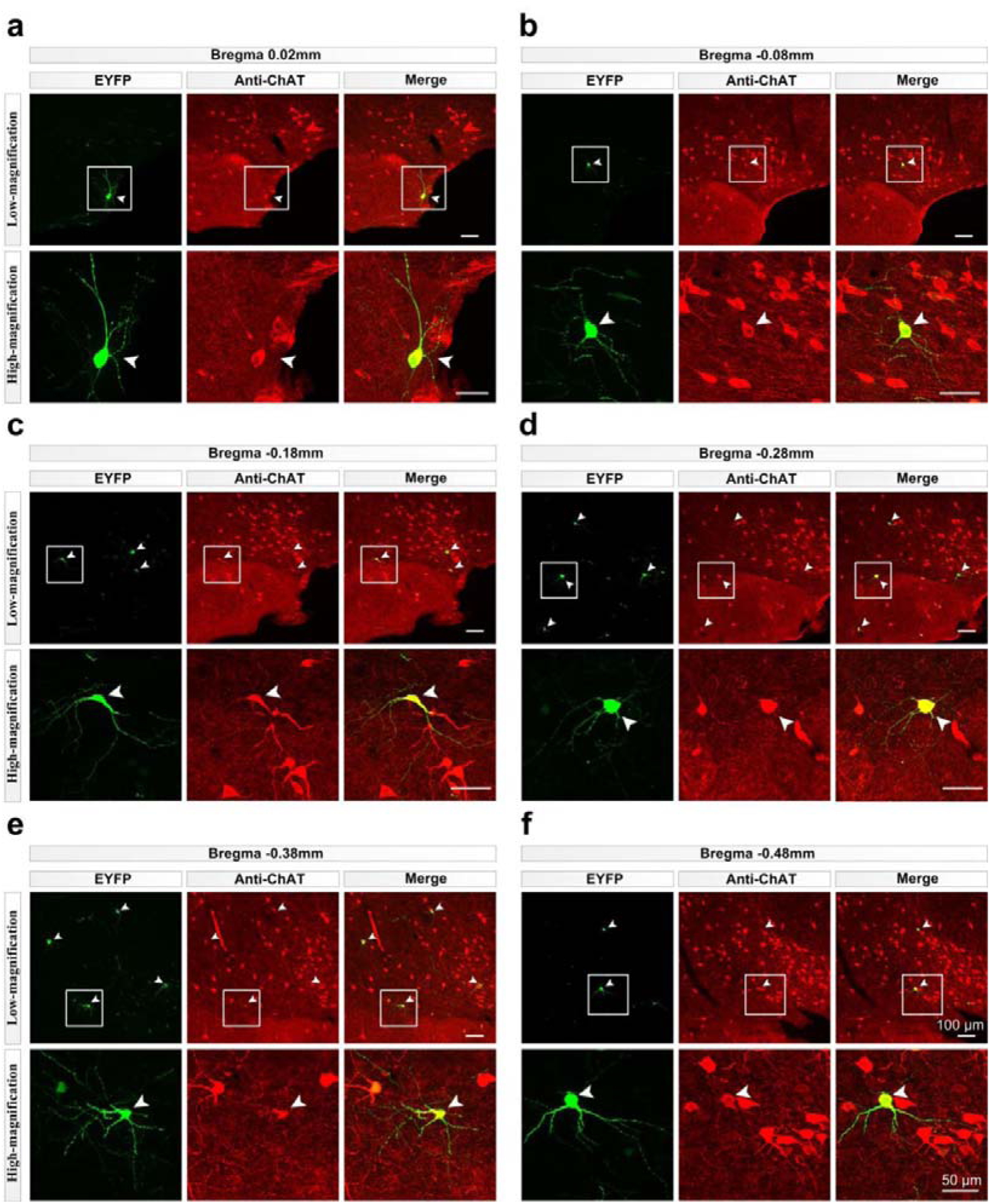
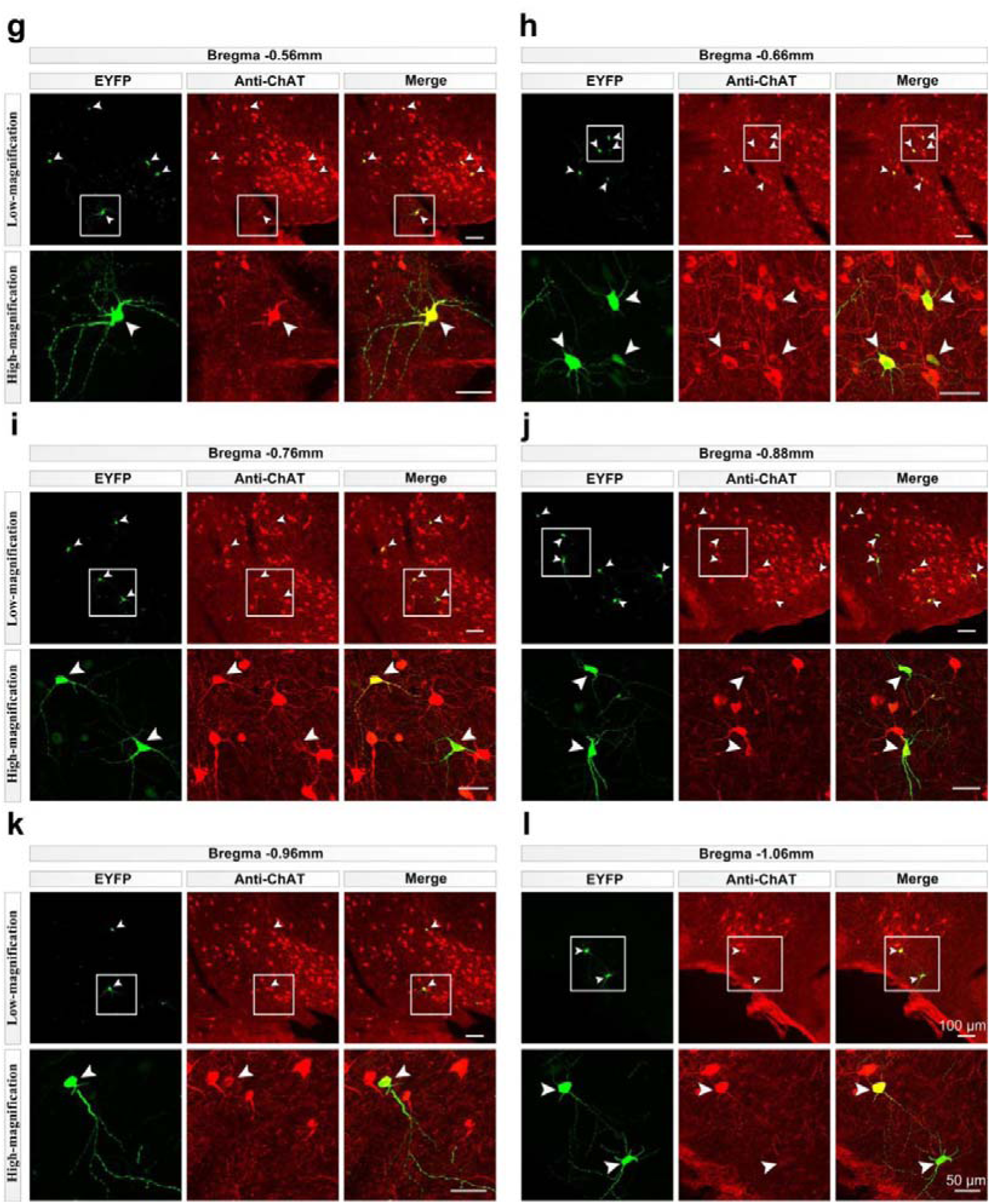
Cell-type-specific, sparse and super-bright BF cholinergic neuron labeling. Twelve 50-µm thickness coronal sections ranging from Bregma 0.02 mm to Bregma - 1.06 mm of the injection site were selected from a cslAAV^20,000^-labeled ChAT-Cre transgenic mouse sample and displayed with a 50-µm intervals and immunostained with anti-ChAT antibody. Upper panels of **a-l** were low-magnification images showing overall labeling on each section and lower panels were high-magnification images of the corresponding boxes. Colabeled neruons were indicated by white arrowheads. This brain sample contained 62 labeled neurons (mouse 4 in **Supplementary Table 1**) and all of them were ChAT-positive and approximately 39 neurons were listed here.

**Supplementary figure 13.**
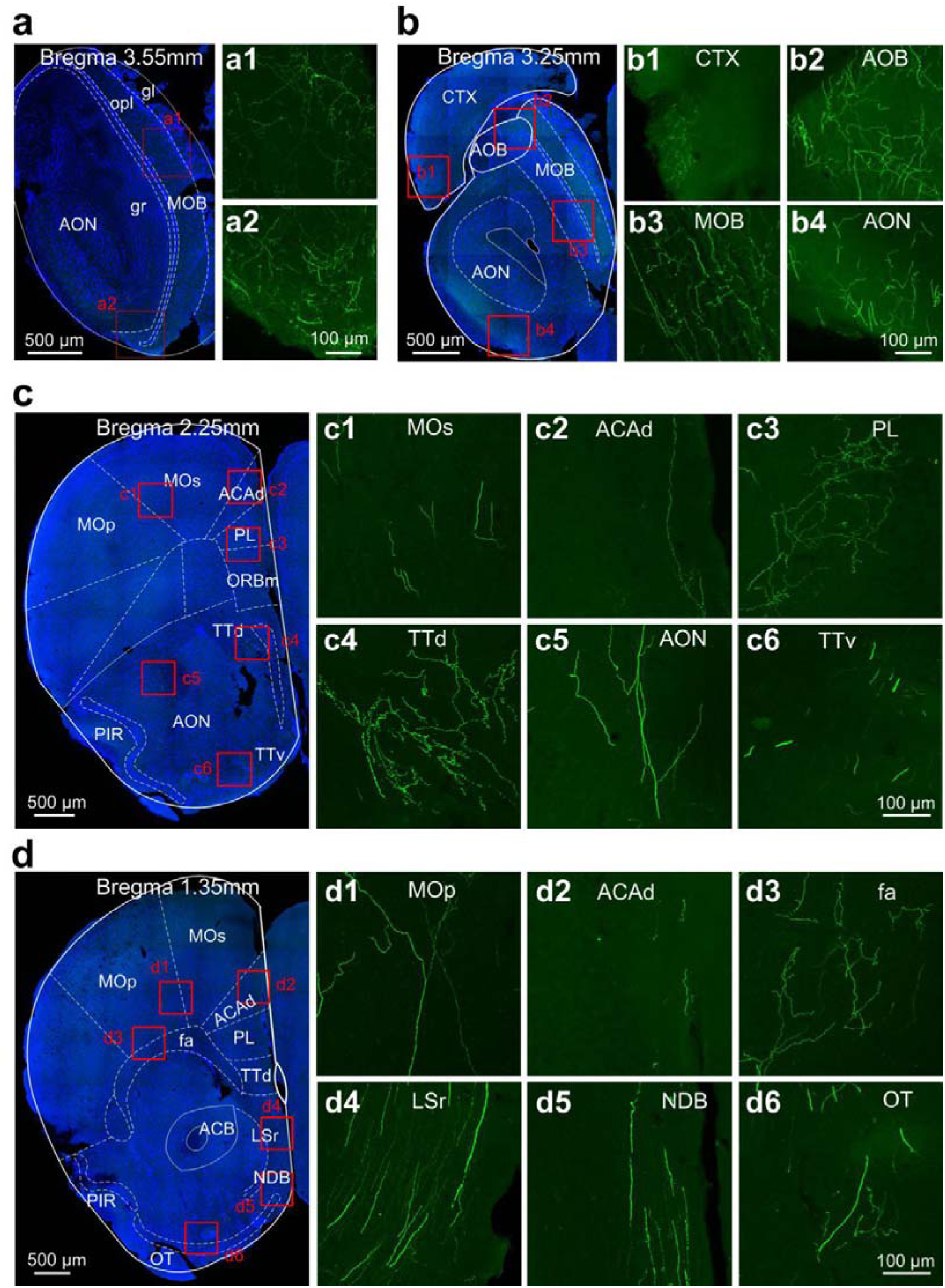
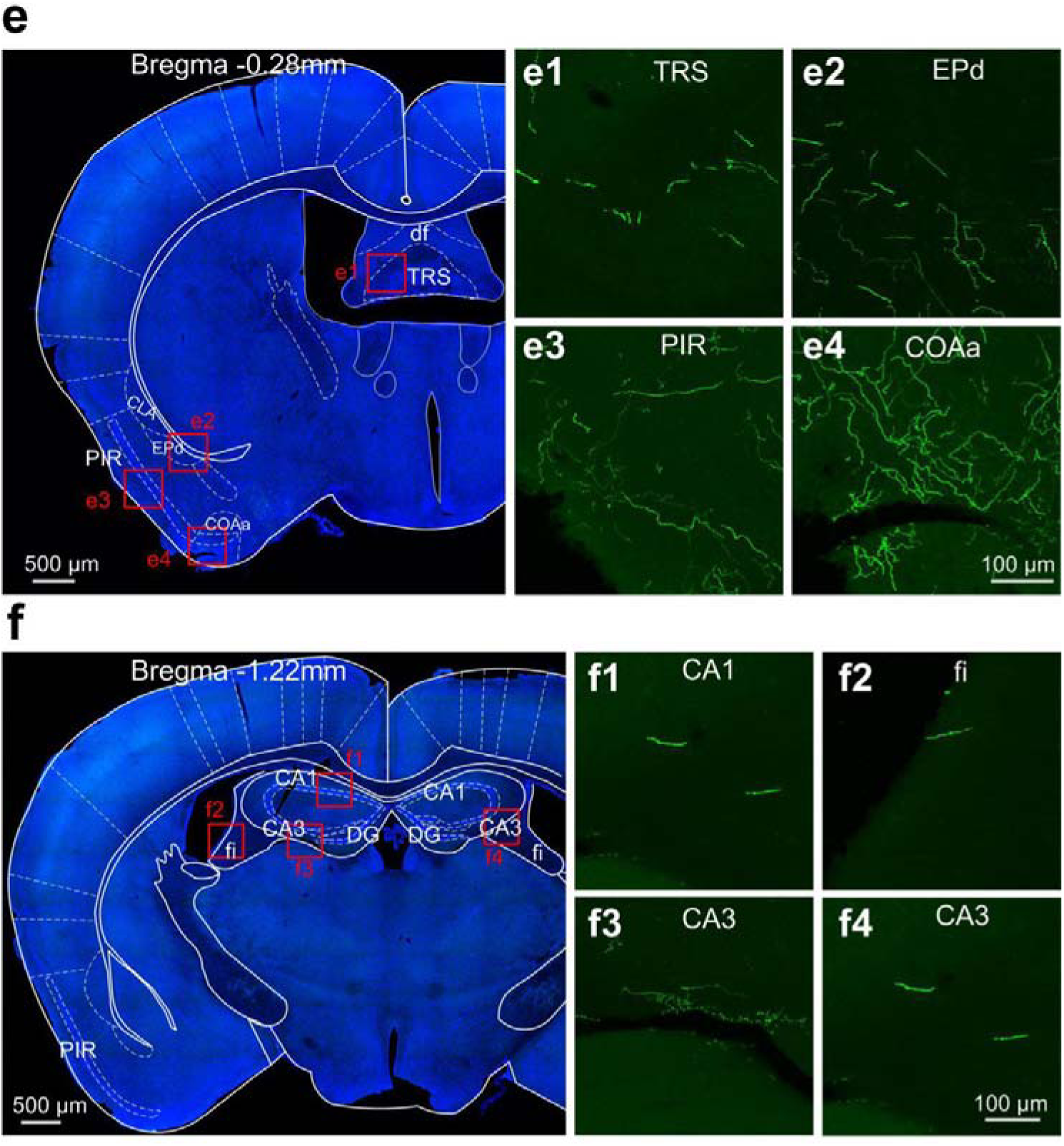
Brain-wide long-range projections of BF cholinergic neurons. Six coronal sections from a representative cslAAV^20,000^-labeled ChAT-Cre transgenic mouse brain sample were selected to show long-range projections of BF cholinergic neuron**s**. **a1-a2**, **b1-b4**, **c1-c6**, **d1-d6**, **e1-e4** and **f1-f4** were maximum-intensity projections of the respective boxed regions in **a-f** showing the details of the axonal projections in different brain regions. Abbreviations: ACAd, Anterior cingulate area, dorsal part; ACB, nucleus accumbens; AOB, accessory olfactory bulb; AON, accessory olfactory nucleus; CA1, field CA1 of hippocampus; CA3, field CA3 of hippocampus; CLA, claustrum; COAa, cortical amygdalar area, anterior part; CTX, cerebral cortex; df, dorsal fornix; DG, dentate gyrus; ENd, endopiriform nucleus, dorsal part; fa, fcorpus callosum, anterior forceps; fi, fimbria of the hippocampus; LSr, lateral septal nucleus, rostral (rostroventral) part; MOB, main olfactory bulb; MOBgl, main olfactory bulb, glomerular layer; MOBgr, main olfactory bulb, granule layer; MOBopl, main olfactory bulb, outer plexiform layer; MOp, primary motor area; MOs, secondary motor area; NDB, diagonal band nucleus; ORBm, orbital area, medial part; OT, olfactory tubercle; PIR, piriform area; PL, Prelimbic area; TRS, triangular nucleus of septum; TTd, taenia tecta, dorsal part; TTv, taenia tecta, ventral part.

**Supplementary figure 14.**
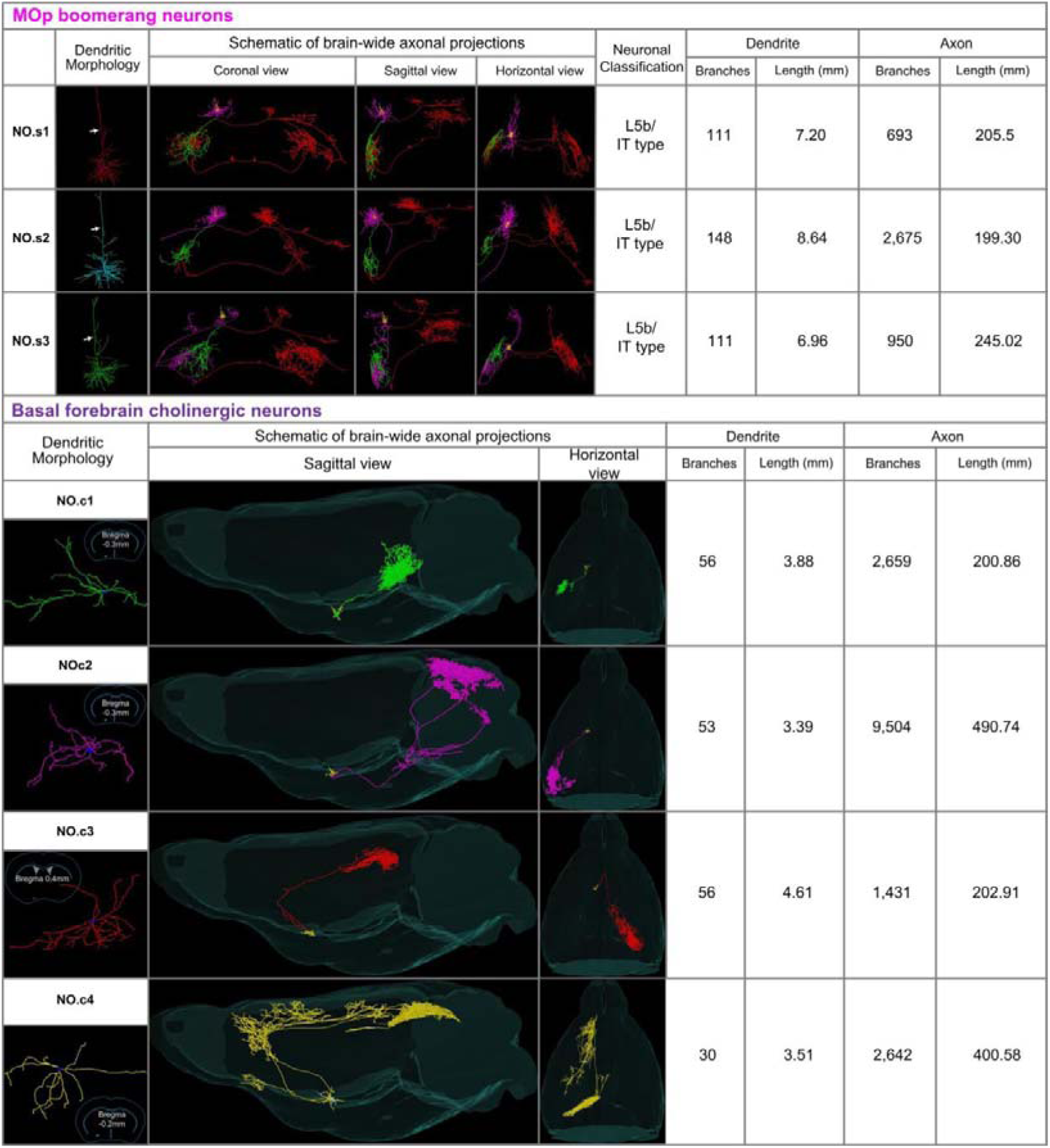
Quantitative analysis of MOp boomerang neurons. (No. **s1-s3**) **and BF cholinergic neurons** (No. **c1-c4**). For three boomerang neurons, dendrites were depicted in yellow, local axons were in magnet, left branches and right branches originated from ipsilateral external capsule were in green and red color, respectively for better differentiation. White arrows indicate apical dendrites. For four cholinergic neurons, coarse outlines and green dots indicated approximate locations of cell body, brain-wide long-range projections were demonstrated in sagittal and horizontal views, dendritic morphologies were depicted in yellow (No. **c1-c3**) and white (No. **c4**).

**Supplementary figure 15.**
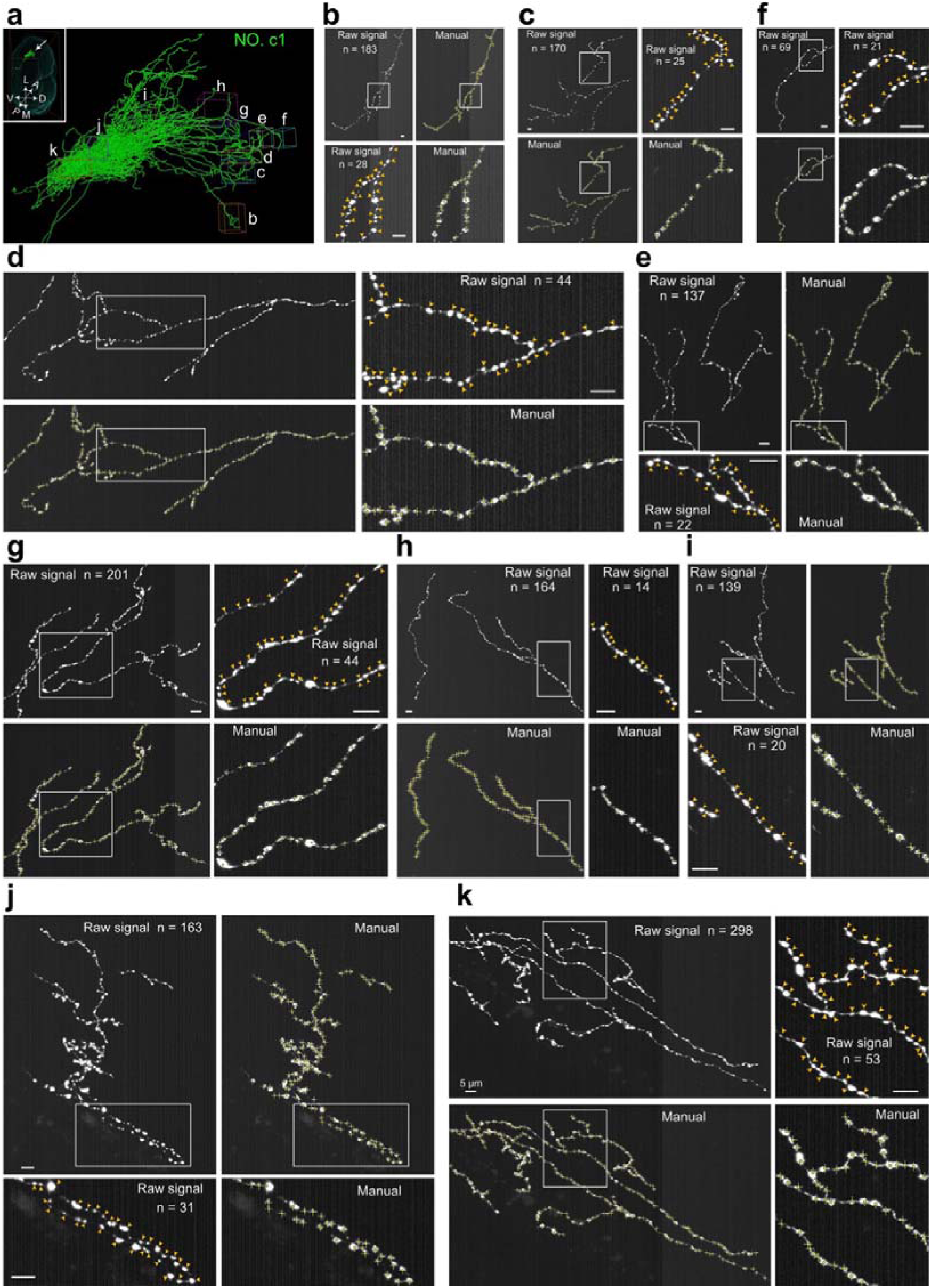
Schematic of identifying presynaptic axonal boutons with a representative cholinergic neuron (No. c1). **(a)** Overview of ten regions of interest (ROIs, **b-k**) randomly selected from No. **c1** for bouton identification. **(b-k)** Demonstration of raw signals (indicated by yellow arrowheads) and manual identification of boutons with ImageJ (indicated by yellow crosses) of ten ROIs in **a**.

**Supplementary Table 1.** Quantifications of starter cells and total input cells of lAAV-DIO-GT/RG and sAAV-DIO-GT/RG labeling groups.

| Total number of starter cells |  |  |  |  |  |  |
| --- | --- | --- | --- | --- | --- | --- |
| | Mouse 1 | Mouse 2 | Mouse 3 | Mouse 4 | Mouse 5 | Mean ± sem |
| IAAV-DIO-GT/RG | 2,999 | 2,308 | 2,396 | 1,940 | 2,286 | 2,386 ± 172 |
| sAAV-DIO-GT/RG | 1,842 | 2,406 | 2,155 | 1,974 | 1,817 | 2,039 ± 127 |
| Total number of input cells across the whole brain |  |  |  |  |  |  |
| IAAV-DIO-GT/RG | 150,915 | 132,027 | 113,278 | 115,100 | 105,647 | 123,393 ± 9,076 |
| sAAV-DIO-GT/RG | 81,771 | 70,309 | 72,182 | 43,507 | 54,665 | 64,487 ± 7,617 |

**Supplementary Table 2.**
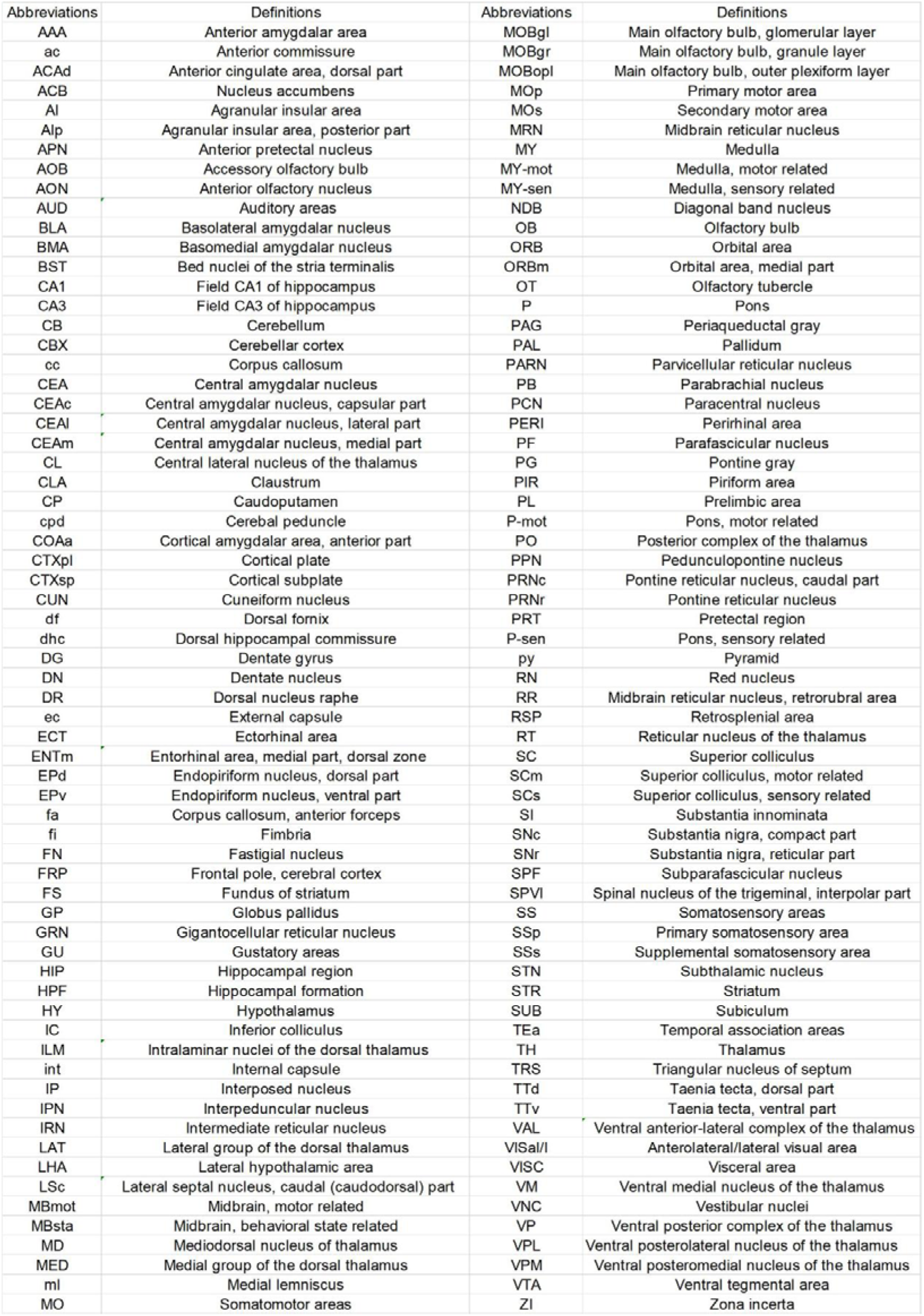
Abbreviations of anatomical structures according to the Allen Brain Atlas (ABA, http://mouse.brain-map.org/).

**Supplementary Table 3.** The total numbers of labeled neurons across the whole brain after labeling with the indicated lAAVs, sAAVs and cslAAVs in different regions.

| Non-cell-type-specific labeling with IAAVs (wild-type mice) |  |  |  |  |  |  |  |
| --- | --- | --- | --- | --- | --- | --- | --- |
| Injection site |  | Coordinates | Cell number across the whole brain |  |  |  |  |
| | | | Mouse 1 | Mouse 2 | Mouse 3 | Mouse 4 | Mean ± sem |
| MOp | IAAV<sup>20,000</sup> | AP= 1.54<br>ML= -1.60<br>DV= -1.60 | 652 | 916 | 688 | 329 | 646 ± 121 |
| | IAAV<sup>200,000</sup> | | 74 | 68 | 58 | 77 | 69 ± 4 |
| | sAAV<sup>20,000</sup> | | 100 | 182 | 260 | 236 | 195 ± 35 |
| | sAAV<sup>40,000</sup> | | 103 | 183 | 151 | 114 | 138 ± 18 |
| | sAAV<sup>80,000</sup> | | 66 | 36 | 16 | 59 | 44 ± 11 |
| IAAV<sup>200,000</sup> |  |  |  |  |  |  |  |
| CA3 | AP= -1.7; ML= -2.0;<br>DV= -2.0 | | 49 | 75 | 59 | | 61 ± 8 |
| CA1 | AP= -1.7; ML= -2.0;<br>DV= -1.5 | | 150 | 98 | 137 | | 128 ± 16 |
| Cell-type-specific labeling with csIAAVs (Cre-driver transgenic mice) |  |  |  |  |  |  |  |
| Mop | csIAAV<sup>20,000</sup> | AP= 1.54<br>ML= -1.6<br>DV= -1.6 | 497 | 573 | 580 | | 550 ± 27 |
| Basal forebrain | csIAAV<sup>20,000</sup> | AP= 0.21<br>ML= -1.5<br>DV= -5.5 | 46 | 58 | 48 | 62 | 54 ± 4 |
| VTA | csIAAV<sup>8,000</sup> | AP= -3.4<br>ML= -0.3<br>DV= -4.1 | 116 | 137 | 159 | 214 | 157 ± 21 |

**Supplementary Table 4.** Quantifications of presynaptic axonal boutons of four cholinergic neurons (No. c1-c4).

|  | No.c1 |  |  | No.c2 |  |  | No.c3 |  |  | No.c4 |  |  |
| --- | --- | --- | --- | --- | --- | --- | --- | --- | --- | --- | --- | --- |
|  | Number of boutons | Axonal length (μm) | density /μm | Number of boutons | Axonal length (μm) | density /μm | Number of boutons | Axonal length (μm) | density /μm | Number of boutons | Axonal length (μm) | density /μm |
| ROI-1 | 183 | 539.7119 | 0.3391 | 59 | 217.998 | 0.2706 | 43 | 154.5866 | 0.2782 | 45 | 142.4557 | 0.3159 |
| ROI-2 | 170 | 652.3893 | 0.2606 | 45 | 146.6952 | 0.3068 | 112 | 336.6172 | 0.3327 | 41 | 108.3174 | 0.3785 |
| ROI-3 | 209 | 615.2699 | 0.3397 | 43 | 156.8266 | 0.2742 | 41 | 178.8266 | 0.2293 | 113 | 383.7817 | 0.2944 |
| ROI-4 | 298 | 919.7885 | 0.324 | 38 | 125.9487 | 0.3017 | 58 | 210.1786 | 0.276 | 46 | 197.7769 | 0.2326 |
| ROI-5 | 69 | 177.9003 | 0.3879 | 39 | 160.4843 | 0.243 | 41 | 142.1778 | 0.2884 | 52 | 189.0938 | 0.275 |
| ROI-6 | 137 | 407.4272 | 0.3363 | 33 | 96.53236 | 0.3419 | 69 | 233.0409 | 0.2961 | 57 | 169.729 | 0.3358 |
| ROI-7 | 201 | 568.9249 | 0.3533 | 59 | 192.9515 | 0.3058 | 46 | 172.0581 | 0.2674 | 64 | 240.632 | 0.266 |
| ROI-8 | 163 | 772.5238 | 0.211 | 69 | 151.3374 | 0.4559 | 62 | 248.3191 | 0.2497 | 43 | 130.3187 | 0.33 |
| ROI-9 | 164 | 490.4556 | 0.3344 | 48 | 176.4415 | 0.272 | 73 | 298.1006 | 0.2449 | 146 | 386.982 | 0.3773 |
| ROI-10 | 139 | 441.6116 | 0.3148 | 36 | 147.8423 | 0.2435 | 56 | 171.7005 | 0.3261 | 34 | 106.6103 | 0.3189 |
| Mean | 173.3 | 558.6003 | 0.3201 | 46.9 | 157.305786 | 0.3015 | 60.1 | 214.5606 | 0.2789 | 64.1 | 205.56975 | 0.3124 |
| SEM | 18.637507 | 64.5331285 | 0.0158 | 3.7340773 | 10.6693277 | 0.0197 | 6.8190094 | 20.31245507 | 0.0106 | 11.485692 | 32.7300585 | 0.0148 |
| Estimated number of boutons | 64,295 ± 3,174 |  |  | 147,958 ± 9,668 |  |  | 56,592 ± 2,151 |  |  | 125,138 ± 5,928 |  |  |

**Supplementary Table 5.**
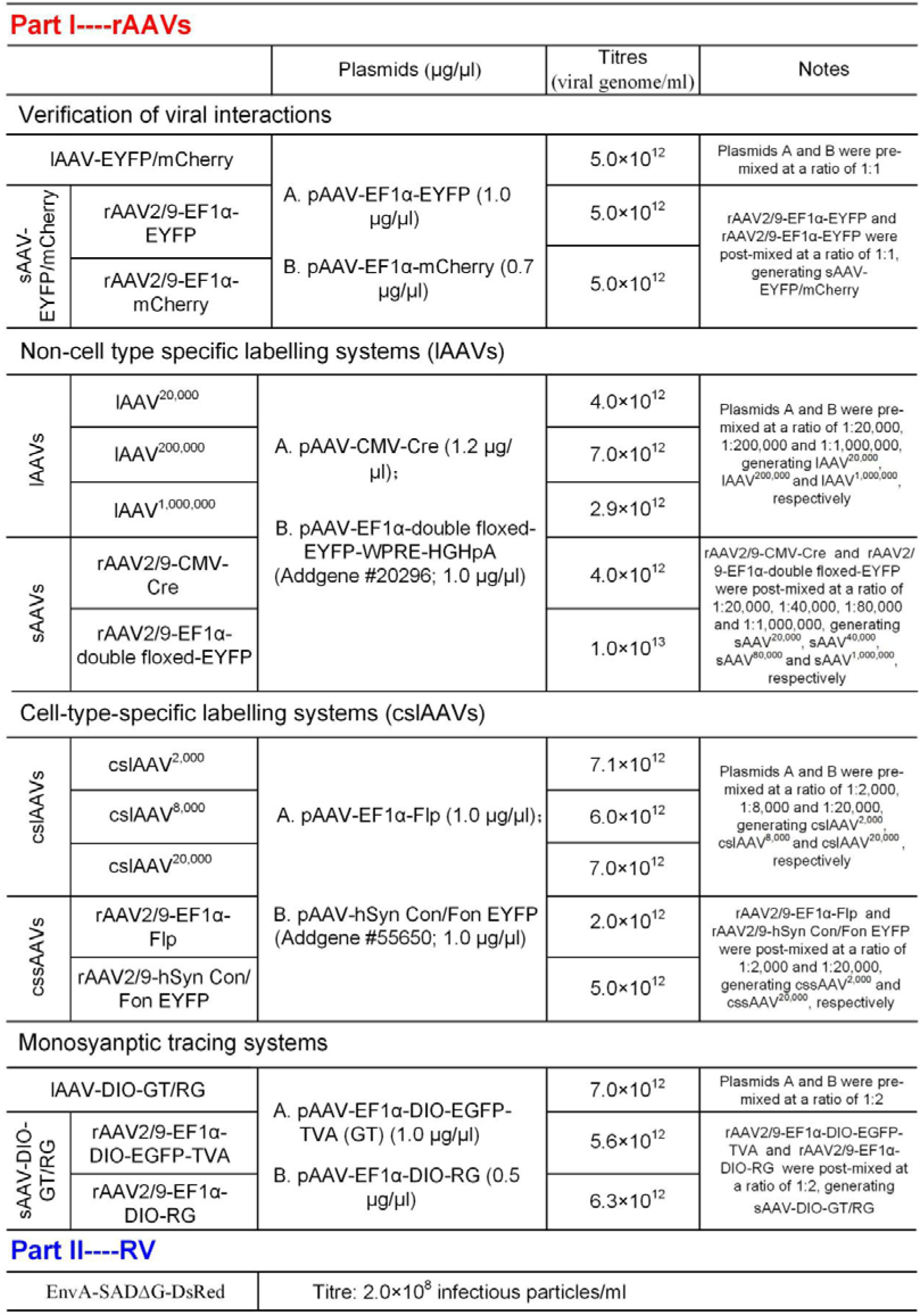
Summary of viruses used in this study.

## Notes

### Competing Interest Statement

The authors have declared no competing interest.

